# De novo design of peptides localizing at the interface of biomolecular condensates

**DOI:** 10.1101/2025.05.09.653111

**Authors:** Timo N. Schneider, Marcos Gil-Garcia, Marco A. Bühler, Lucas F. Santos, Lenka Faltova, Gonzalo Guillén-Gosálbez, Paolo Arosio

## Abstract

The interface of biomolecular condensates has been shown to play an important role in processes such as protein aggregation and biochemical reactions. Targeted modulation of these interfaces could, therefore, serve as an effective strategy for engineering condensates and modifying aberrant behaviors. However, the molecular grammar driving the preferential localization of molecules at condensate interfaces remains largely unknown. In this study, we developed a computational pipeline that combines highthroughput coarse-grained simulations, machine learning, and mixed-integer linear programming to design peptides that selectively partition at the interfaces of specific condensate targets. Using this workflow, we designed and synthesized peptides that localize at the interface of three distinct condensates formed by different intrinsically disordered protein regions (IDRs). These peptides exhibit surfactant-like architectures, with one tail incorporated into the condensate and the other excluded from the dense phase. In all cases, the tail entering the condensates is enriched in aromatic residues, while the sequence of the excluded tail varies among the IDRs. For hnRNPA1- and LAF1-IDRs, the excluded tail is enriched in lysines and matches the net charge of the condensate-forming protein, promoting electrostatic repulsion. In the case of DDX4-IDR, which exhibits the lowest charge density, the excluded tail mainly consists of uncharged valine residues, which exhibit negligible interactions with the scaffold protein. These results highlight the importance of the net charge of the scaffold as a key physicochemical parameter for designing peptides with preferential interfacial localization. Overall, our pipeline represents a promising strategy for the rational design of interface-localizing peptides and the identification of the corresponding molecular grammar.

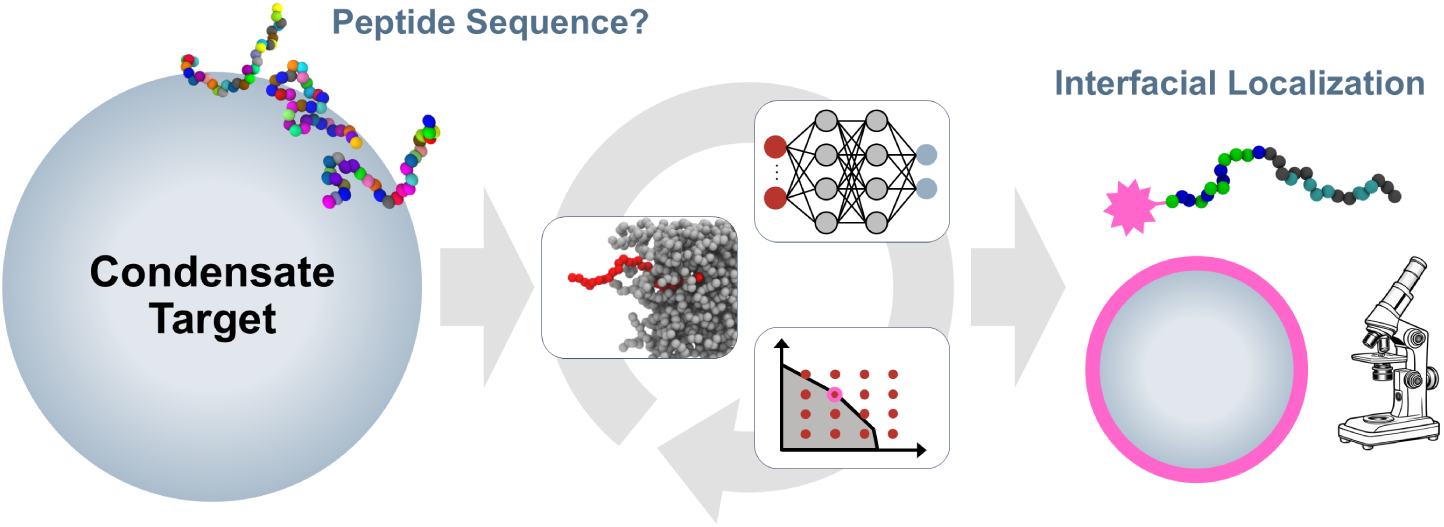

## Introduction

Growing evidence shows that cells regulate biochemical reactions in space and time through the formation of membraneless organelles, also known as biomolecular condensates.^1–4^ These protein- and RNA-rich condensates have been linked to several cellular functions—such as RNA metabolism and stress responses^1^—as well as disease-associated dysfunctions, including progression to arrested states and formation of amyloid fibrils.^5,6^ In addition to changes in local concentrations, condensates can affect biochemical functions by other emergent properties at the mesoscale. One of these properties which is attracting increasing attention is the interface between the dense and dilute phase. This interface has been shown to promote the formation of disease-associated fibrils of hnRNPA1,^7,8^ as well as liquid-to-solid transition of FUS.^9^ Condensate interfaces can also promote the aggregation of client proteins, as shown with mutant huntingtin polyglutamine in vivo, ^10^ and also *α*-synuclein in vitro.^11,12^ Moreover, interfaces can define electric potentials and promote redox reactions. ^13–15^

Targeted modulation of the interface of condensates, without perturbing their interior and other mesoscale properties, could therefore be an attractive strategy for engineering condensates and correcting aberrant behaviors. A traditional strategy in colloid science for altering interfacial properties involves the use of surfactant molecules. However, this approach presents challenges in the context of condensates, as the dilute and dense phases are more similar to each other than, for example, in oil-in-water emulsions.^16,17^ Both phases contain substantial amounts of water and dissolved ions, with the volume fraction of water in the dense phase estimated to be around 70%.^18–21^ Therefore, it is more challenging to design surfactant-like molecules that would preferentially partition at the interface of biomolecular condensates. Furthermore, interfacial localization is expected to strongly depend on the identity of the condensate. Currently, only a few proteins have been shown to exhibit preferential interfacial partitioning,^7,10–12,22–25^ and some condensates exhibit colloidal stability based on mechanisms reminiscent of Pickering emulsions.^26^ However, to our knowledge preferential interfacial localization has not been demonstrated for small molecules or peptides. Moreover, a high-throughput experimental strategy for extensive screening is still lacking. As a result, the guiding principles for preferential localization at the interface of biomolecular condensates remain elusive.

The predictive capability of molecular dynamics simulations using coarse-grained force fields^27–30^ has shown promising results for designing polymers, peptides, and disordered proteins for a variety of applications.^31–34^ The efficiency of the design process can be improved with machine learning models, enabling active learning in sequence space. The inverse design problem, i.e. selecting new sequences from a large space, guided by a trained model, is commonly tackled with genetic algorithms or by performing local optimization in a continuous, potentially learned representation of the sequence space.^35–42^ One drawback of these approaches is the risk of getting trapped in local optima due to the nonconvex nature of the optimization model. Recent work^43,44^ has enabled the identification of globally optimal inputs for trained neural networks using mixed-integer linear programming (MILP), which can be used to mimic the behaviour of complex systems, including nonconvex models based on first principles.

Here we have developed a computational pipeline based on a combination of coarsegrained molecular simulations, machine learning and MILP for the de novo design of peptides partitioning at the interface of biomolecular condensates. Specifically, coarse-grained molecular simulations were integrated with an active learning algorithm involving the training and optimization of a surrogate model based on a neural network. The optimization problem containing the trained neural network was formulated as a mixed-integer linear programming formulation, which led to globally optimal sequences for each iteration. We applied this framework to de novo design and experimentally validate peptides which partition at the interface of condensates formed by the intrinsically disordered domains of various proteins associated with biological condensates, such as hnRNPA1, LAF-1, and DDX4. The designed peptides accumulated at the protein interface *in vitro*, as intended, and displayed surfactant-like architectures, with each case exhibiting a distinct composition. Based on these results, we identified design rules for interface-partitioning peptides, highlighting the net charge of the condensate-forming protein as a key physicochemical feature.

## Results and Discussion

### Overall workflow

We chose to design peptides comprising 30 amino acid residues to achieve a balance between sequence variability and accessibility for chemical synthesis. The de novo design of such peptides encompasses selecting optimal sequences of amino acids from a vast design space. To address this challenge, we therefore combined high-throughput coarse-grained simulations with an active learning algorithm for a more efficient navigation. A representation of the entire workflow is shown in Figure 1.

**Figure 1:**
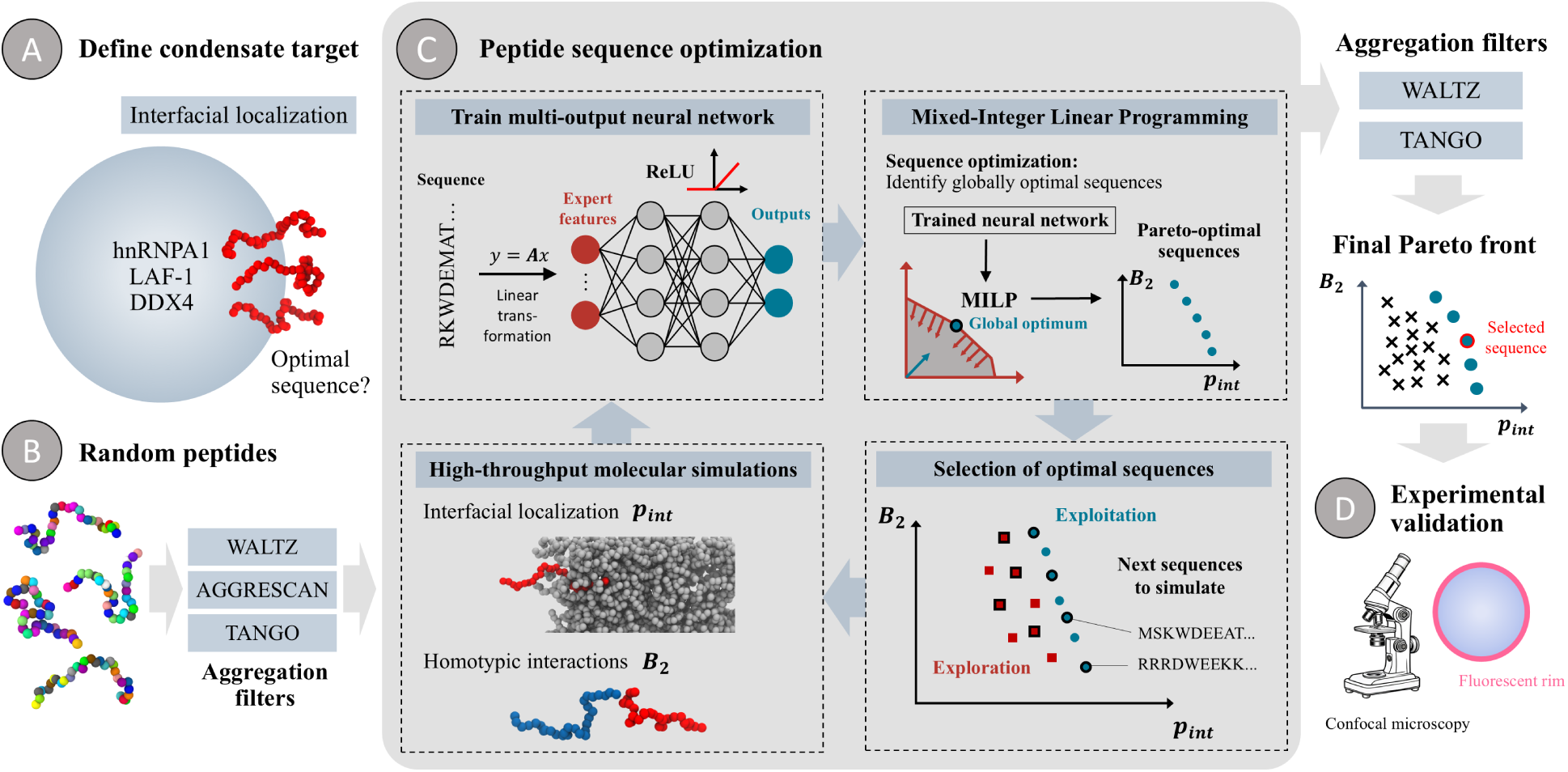
Overall workflow for designing peptides partitioning at the interface of biomolecular condensates: After definition of a condensate target (A) randomly generated peptides were screened to exclude aggregation-prone regions (B) and entered the optimization cycle (C). Interface partitioning and homotypic interactions were quantified using coarse-grained molecular simulations, and the resulting data was used to train a surrogate model (neural network). In a next step, we leveraged mixed-integer linear programming to identify globally optimal sequences, leading to new peptides to simulate. Upon convergence of this optimization cycle, we experimentally validated our predictions using confocal microscopy (D).

After defining the condensate target (Figure 1A), the analysis began with a set of fully randomized peptides, in which each of the 20 amino acids was uniformly sampled. To enhance the synthesizability and stability of the designed peptides, we implemented a filter to prevent the emergence of any aggregation-prone regions. Specifically, we applied three different sequence-based aggregation predictors: Waltz,^45^ TANGO,^46^ and AGGRESCAN^47^ (Figure 1B). This strategy led to 300 initial sequences, without introducing any bias towards existing biological protein sequences.

Next, the sequences were introduced into the optimization loop (Figure 1C), where we aimed at simultaneously maximizing the interface partitioning of the single peptide and minimizing homotypic interactions. This is because strong self-association may lead to aggregation, reducing partitioning at the interface at higher concentrations. Moreover, the minimization of homotypic interactions serves as an additional safeguard against generating highly aggregation-prone or insoluble sequences that may not be identified by existing aggregation predictors. We performed two separate coarse-grained simulations for each peptide: in the first simulation, we quantified the probability of a single peptide to be localized at the interface of a minimalistic condensate formed by the protein of interest (*p*_int_); in the second simulation, we included two copies of the peptide to measure homotypic interactions, quantified by the second virial coefficient (*B*_2_).

In the next step, we trained a surrogate model (multi-output neural network) on the simulation results to predict both interface partitioning as well as homotypic interactions. The network uses engineered features, obtained through a linear transformation of the peptide sequence, as inputs, and ReLU as the activation function. Both aspects were crucial for the next step: the bi-objective optimization problem with the trained neural network embedded was reformulated as a MILP problem^43,44^ and solved to maximize simultaneously *p*_int_ and *B*_2_. This led to Pareto solutions^48^—a set of globally optimal, non-dominated sequences that cannot improve one objective function without compromising the other one—with mathe-matical guarantees within the massive sequence design space of size 20^30^. In contrast with, for instance, genetic algorithms, this approach avoids stagnation in local optima, enabling the construction of the true Pareto front of the surrogate model within the desired optimality gap. Combined with an exploration strategy, this process typically selected 100 new sequences for further simulations, restarting the optimization cycle. Following convergence of the cycle, we applied final aggregation filters and selected a sequence for experimental validation (Figure 1D). A fluorescently tagged variant of the peptide was then synthesized, and its localization within the condensate system was analyzed *in vitro* using confocal microscopy.

### Simulation, machine learning, and optimization setup

Although the molecular simulations were performed at a coarse-grained level, computational requirements were still significant. Simulations were therefore designed to balance accuracy and throughput. We chose the Mpipi force field,^28^ a one-bead per residue model, due to its physics-based parametrization strategy and capability of explaining experimental data of disordered proteins. We developed a custom simulation procedure to quantify the interfacial localization of peptides, modeling the condensate as a minimalistic slab containing 16 protein copies. Using the adaptive biasing force method,^49,50^ we mapped the potential of mean force (PMF) as a function of peptide position—dilute, interface, or dense phase—as illustrated in Figure 2A. To accelerate convergence, we employed a stratification strategy (Materials and Methods), which also enabled the inclusion of two peptides per simulation without interaction, further increasing simulation throughput. The free energy differences between the dense phase and the interface (Δ*G*_1_) and between the dense and dilute phases (Δ*G*_2_) were extracted from the PMFs and used to compute 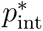, our objective for interface partitioning. 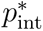 can be approximated as proportional to the probability that the molecule localizes at the interface (*p*_interface_) relative to the bulk of the dilute and dense phase (a detailed derivation is provided in the Supplementary Information):

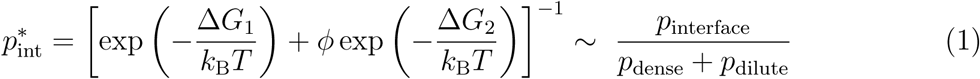

**Figure 2:**
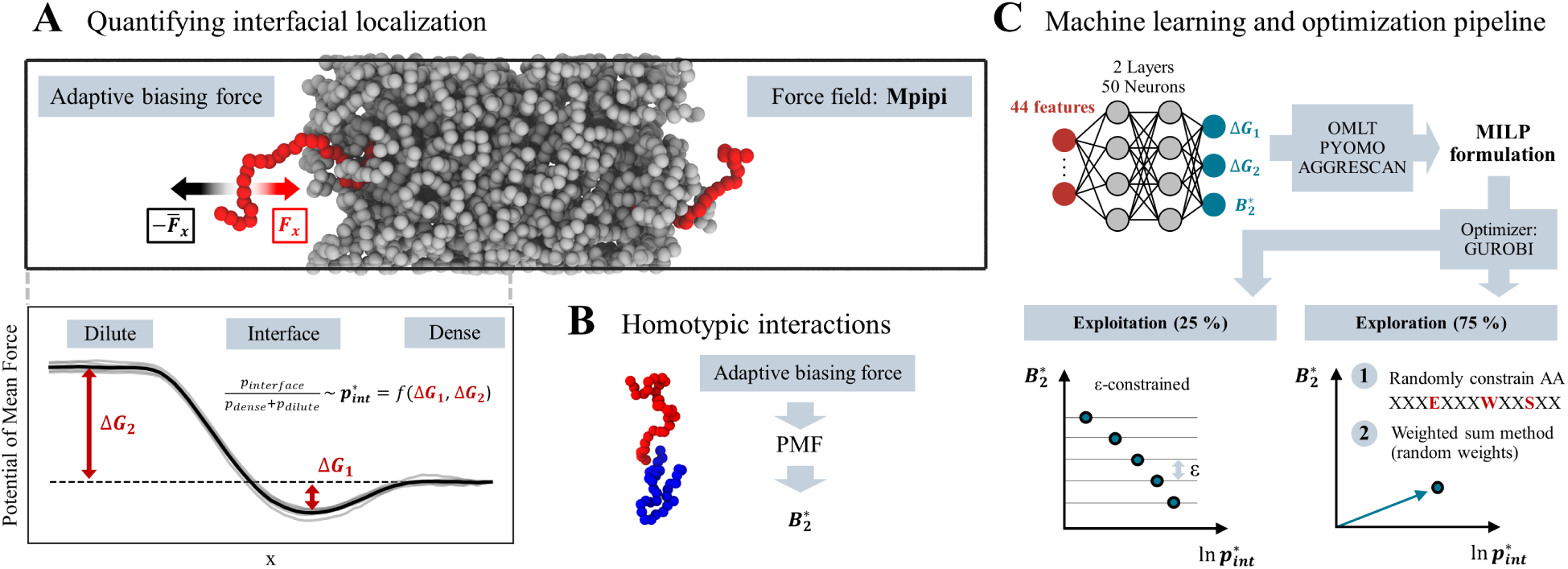
Molecular simulations utilizing the coarse-grained force field Mpipi, in conjunction with the adaptive biasing force method, were employed to quantify interfacial localization (A) and homotypic interactions (B). An ideal peptide demonstrates high interface parti-tioning (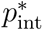), which corresponds to high values of both Δ*G*_1_ and Δ*G*_2_, while exhibitingself-repulsion, reflected by a high second virial coefficient *B*^∗^_2_. A multi-output neural network was trained on the simulation results (C) and translated into a mixed-integer linear programming formulation, which was then optimized with distinct strategies for exploration and exploitation.

Here, *ϕ* represents the volume ratio between the dilute and dense phases. A peptide with high interface partitioning would exhibit large values of both Δ*G*_1_ and Δ*G*_2_.

To quantify homotypic interactions, we also used the adaptive biasing force method in a separate simulation to construct the potential of mean force between two peptide copies, employing the center of mass distance as the collective variable (Figure 2B). The PMF was then converted into the second virial coefficient *B*_2_, a key parameter directly linked to homotypic interactions and also phase separation behavior. ^40,51^ To address the highly skewed distribution of values, we applied a bi-symmetric log transformation^52^ finally resulting in the second objective *B*_2_^∗^. Positive values of *B*_2_^∗^ indicate net repulsion, whereas negative values suggest attraction and potential colocalization of the peptide with itself.

In order to identify sequences that maximize both 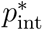 and *B*^∗^_2_, in a next step we trained a surrogate model predicting simulation outcomes (Figure 2C). For each design case, we typically had simulation data only for a few hundred peptide sequences. This small number prevents training a predictor directly on sequence. Instead, we used descriptors, which has been shown to often outperform end-to-end representation learning.^37,53^ We engineered a set of 44 features based on overall composition as well as spatial distribution of charged, aromatic, and strongly self-interacting residues. Additionally, we included features connected to the sequence charge and hydropathy decoration, which play a pivotal role in governing the behavior of disordered proteins.^54,55^ It is important to emphasize that all features can be derived as linear transformations of the one-hot encoded amino acid sequence, which was crucial for later steps. Based on the initial 300 simulations for our first condensate target, which was the disordered domain of hnRNPA1, we compared the predictive performance of multiple models to predict Δ*G*_1_, Δ*G*_2_ and *B∗* (Figure S1). We concluded that a multi-output neural network outperformed elastic net, support vector machine, and gradient-boosted decision tree models. Two hidden layers of 50 neurons each were chosen because larger networks did not provide further improvements in predictive performance (Figure S1).

To address the inverse design problem of identifying optimal sequences, we employed OMLT^44^ to embed the trained neural network into the algebraic modeling framework Pyomo.^56,57^ The piecewise linear nature of the ReLU activation function allowed it to be incorporated into a MILP problem using the big-M reformulation.^43^ To linearize equation (1), which converts the neural network outputs Δ*G*_1_ and Δ*G*_2_ into the objective 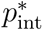, we employed supporting hyperplanes (Supplementary Information). Together, these steps enabled the formulation of the final MILP optimization problem, linking the amino acid sequence inputs to the two objectives via the trained surrogate model. Furthermore, we also used the big-M reformulation to integrate the AGGRESCAN predictor as a constraint, avoiding the emergence of any aggregation-prone regions. The other, more complicated predictors (Waltz, TANGO) were only applied at the end of the optimization cycle, as illustrated in Figure 1. The MILP problem could be solved to global optimality using the optimizer Gurobi.^58^ We utilized the *ε*-constrained method^48^ to generate a Pareto front comprising 25 optimal sequences for the next round of simulations (Figure 2C). In the various iterations it is important to balance exploitation with exploration. To achieve this, 75 sequences per iteration were allocated to exploration, resulting in a total of 100 sequences per iteration. For each exploration sequence, we selected between 2 and 20 positions in the peptide sequence at random and constrained them to randomly chosen amino acids. We then optimized the remaining positions using a weighted-sum approach with randomly generated weights for the two objectives, thereby removing the computational burden of computing *ε*-constraints for each exploration point. This approach aimed to strike a balance between promoting exploration and sampling potentially relevant regions of the sequence space. The entire optimization loop was stopped as soon as the hypervolume of the Pareto front stagnated.

### Peptides targeting the interface of hnRNPA1-LCD condensates

We first applied our approach to design peptides targeting the interface of condensates formed by the low complexity domain (LCD) of hnRNPA1. The optimization pipeline successfully identified peptides exhibiting high interface partitioning and low homotypic interaction, as quantified by coarse-grained simulations. The primary objective 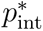 , which measures interface partitioning, improved by approximately three orders of magnitude compared to the initialization (Figure 3A). The algorithm achieved substantial advancement of the Pareto front in the first two iterations, after which the hypervolume plateaued. Optimization was stopped after seven iterations due to stagnation in hypervolume and the similarity of newly suggested sequences to those already simulated. We also compared the construction of the Pareto front using our MILP-based approach with the commonly used genetic algorithm NSGA-II^59^ for the first two iterations. Despite converging on hypervolume, the GA can fail to capture a substantial portion of the true Pareto front (Figure S5). Furthermore, using MILP provides a formal guarantee of convergence to the global optimum of the optimization problem with the ANN embedded, eliminating any uncertainty that can arise with a GA. This highlights the advantage of employing MILP over GA for optimizing the trained neural network.

**Figure 3:**
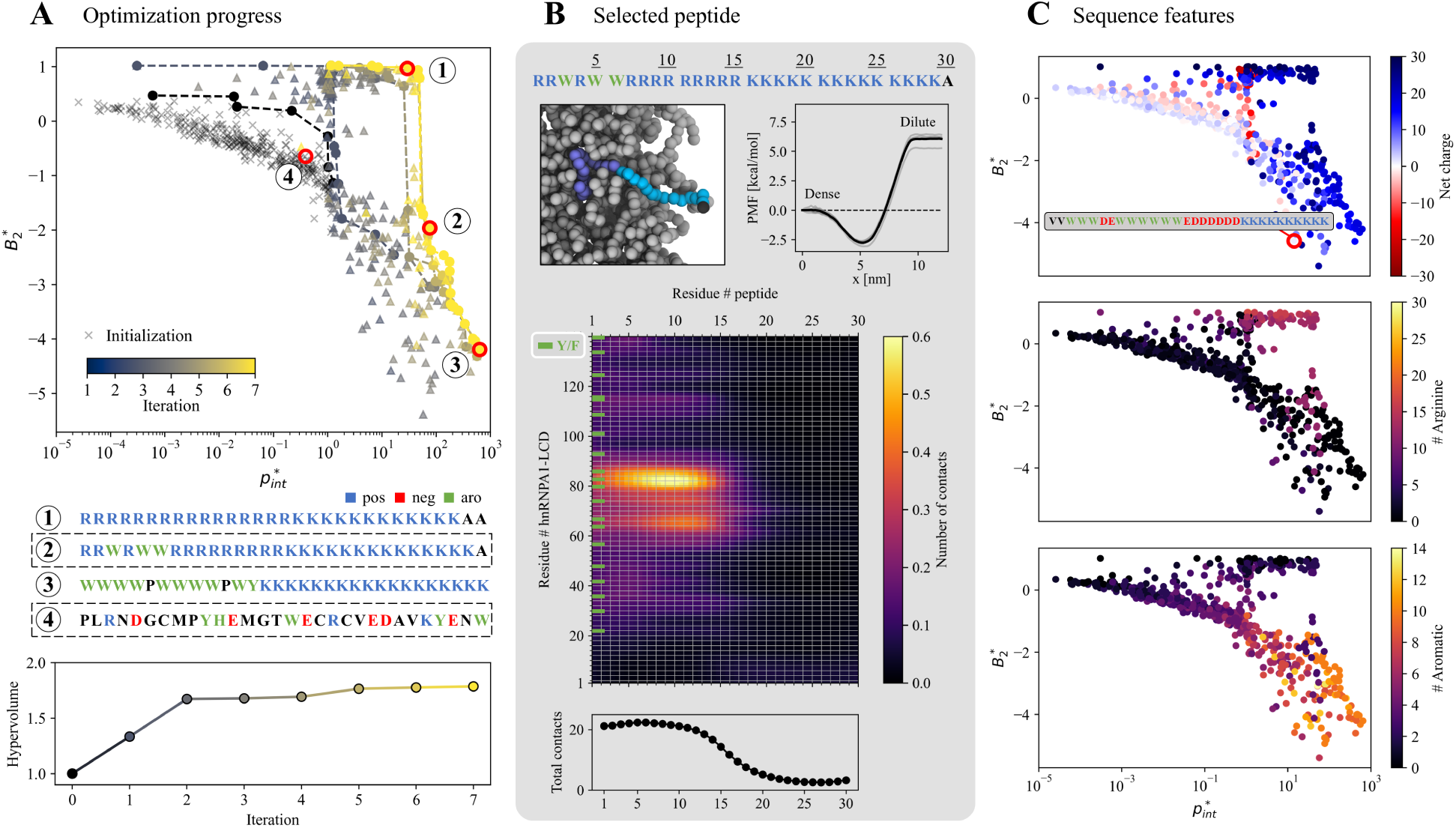
Peptide sequence optimization with hnRNPA1-LCD as the condensate target. (A) Over seven iterations, the primary objective 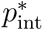, which quantifies interface partitioning, improved by approximately three orders of magnitude. Peptide 2 was later selected for experimental validation of interfacial partitioning, with peptide 4 serving as a control. (B) The selected peptide exhibits surfactant-like behavior: its arginine- and tryptophan-rich end engages in favorable *π*-*π* and cation-*π* interactions with hnRNPA1-LCD, while the positively charged polylysine tail is preferentially excluded from the also positively charged condensate phase. (C) All Pareto-optimal sequences exhibit a high net positive charge, which the algorithm identified as superior to more neutral variants containing aspartic or glutamic acid. Along the Pareto front, aromatic content increases while arginine content decreases with increasing homotypic interactions (low *B*^∗^_2_).

As optimization progresses, the Pareto-optimal peptide sequences became increasingly homogeneous in composition, containing only a subset of the 20 amino acids. This trend is quantitatively reflected in the decreasing average Shannon entropy of the optimal sequences (Figure S11). When sampling sequences from the Pareto front (Figure 3A), it is visible that they exhibit a surfactant-like architecture, as well as a positive net charge. One tail of the peptides is typically enriched in tryptophans and arginines while the other mainly consists of lysines. We selected a peptide from the center of the Pareto front for a closer investigation of the mechanism behind the interfacial localization (Figure 3B). As indicated by the potential of mean force (PMF), this peptide preferentially localizes in the dense rather than the dilute phase, and exhibits even greater affinity for the interface, with the free energy minimum about 2.5 kcal/mol lower than the value in the dense phase. An unbiased simulation (Materials and Methods) was run to analyze the conformational ensemble of the peptide and the per residue contacts with hnRNPA1-LCD. It is visible that the tail rich in arginines and tryptophans heavily interacts with the dense phase formed by the disordered proteins, while the polylysine end remains largely outside the condensate, leading to overall low contact probabilities. This behavior is physically reasonable, as both tryptophan and arginine engage in strong *π*-*π* and cation-*π* interactions, which are known drivers of protein phase separation, including in hnRNPA1-LCD.^60–62^ In fact, the strongest interactions, as identified by the intermolecular contact map, occur with aromatic-rich segments of hnRNPA1 (Figure 3B). By contrast, cation–*π* interactions involving lysine are weaker than those with arginine, a distinction reflected in the Mpipi force field.^28^ Moreover, hnRNPA1-LCD has a net charge of +8, matching the positive charge of the peptide and thereby promoting the exclusion of the polylysine tail from the condensate.

This positive net charge remains a consistent sequence feature across the entire Pareto front (Figure 3C). Peptides with interaction-prone tails containing negatively charged residues, such as aspartic and glutamic acid, were also explored but proved less effective in maximizing the objective functions and were consequently dropped by the algorithm in later iterations. Along the Pareto front, as interface partitioning increases (high 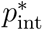, low *B*^∗^_2_), the number of arginines in the sequence decreases while the aromatic fraction increases. At the highest 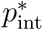 values, sequences no longer contain arginines but are primarily composed of tryptophans and lysines. Aromatic patches are interrupted by prolines (Figure 3A), a consequence of the AGGRESCAN constraint, which prevents the emergence of aggregation-prone regions by incorporating proline as a beta-sheet-breaking residue. ^63^ Notably, Figure 3 shows the direct output of the optimization procedure before the Waltz and TANGO aggregation filters were applied. Once these additional filters are imposed, many peptides with high aromatic content were excluded (Figure S10). Consequently, we selected sequences 2 and 4 (Figure 3A) for experimental validation, as both passed the additional aggregation filters and exhibit different levels of predicted interface partitioning.

We conjugated the synthesized peptides with the fluorescent dye Cy5, before incubation with hnRNPA1-LCD condensates at a protein-to-peptide ratio of 10:1. Analysis by fluorescence confocal microscopy confirmed that peptide 2 exhibits the strong predicted interface partitioning (Figure 4A). Formation of a fluorescent rim was consistent over 5 hours of incubation (Figure S7), and independent of the order in which protein and peptide were added, ruling out mass transfer limitations and confirming that the interfacial localization is thermodynamically driven. In contrast, the control peptide (peptide 4) with a much lower value for 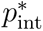 distributed uniformly throughout the dense phase without forming a fluorescent rim (Figure 4B). Interestingly, consistent with its interfacial localization, peptide 2 also caused a significant decrease in average condensate diameter (from 5.6 ± 2.3 *µ*m to 1.7 ± 0.7 *µ*m, mean ± standard deviation, n = 50), as analysed by bright field microscopy (Figure 4C). Dynamic light scattering analysis indicated that peptide 2 also promoted formation of nanometer-sized clusters which are below the limit of detection of microsopy (Figure S9). The addition of control peptide 4 did not significantly affect the condensate diameter, which remained nearly unchanged at 5.5 ± 1.4 *µ*m.

**Figure 4:**
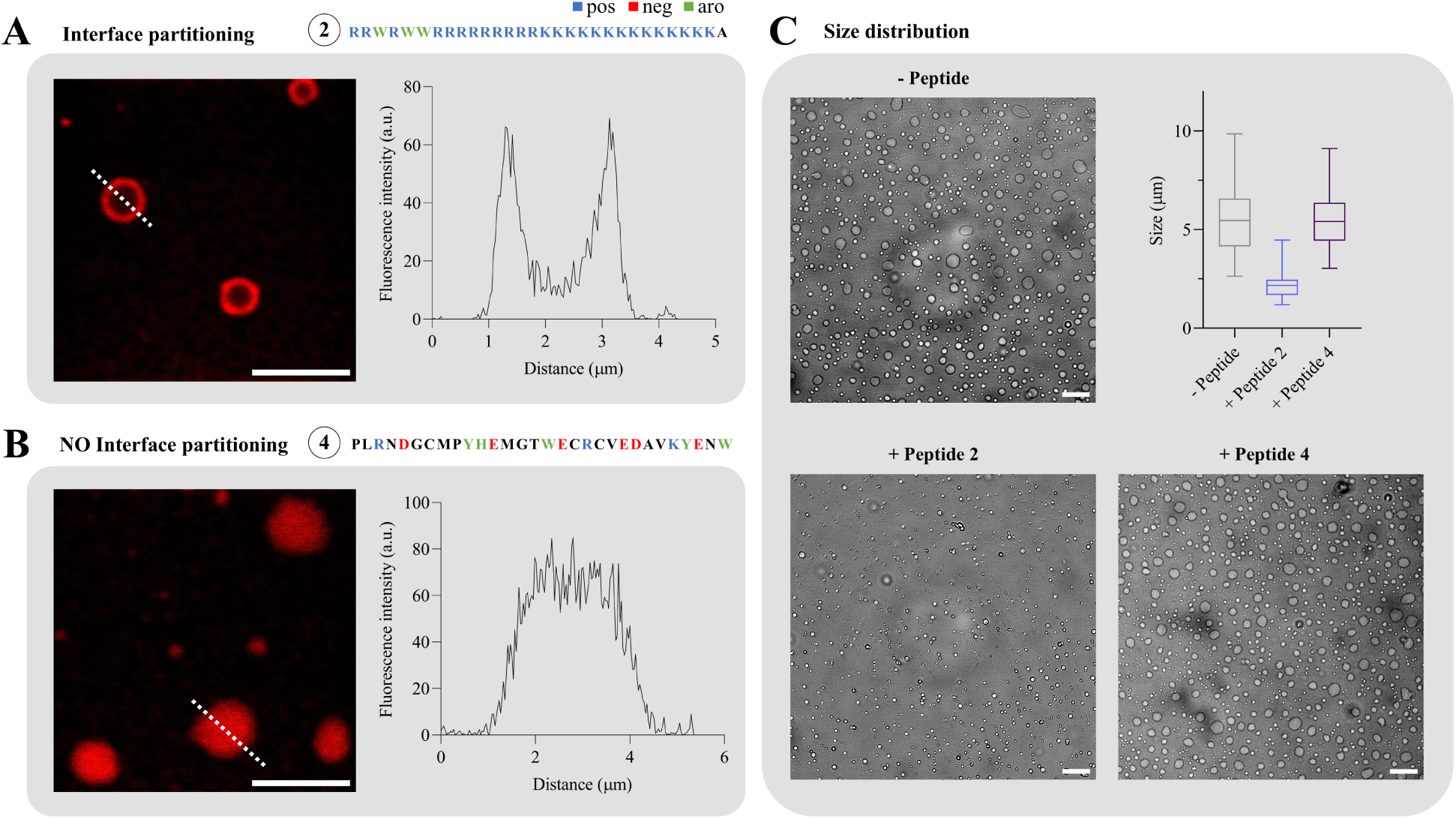
Experimental validation of interface partitioning at hnRNPA1-LCD condensates. Representative confocal microscopy images of the Cy5-labeled (A) selected peptide, with a high predicted propensity to localize at the interface, and (B) control peptide, which is not expected to localize at the interface. The fluorescent rim in (A) confirms the interfacial localization of the selected peptide as predicted by simulations. The amino acid sequences of both peptides are shown. Scale bar: 5 *µ*m. (C) Analyzing size distribution using bright field microscopy reveals that addition of peptide 2 also leads to a decrease in condensate diameter, while addition of peptide 4 does not have a significant impact. Scale bar: 20 *µ*m.

### Designing peptides targeting other protein condensates

We next applied our approach to other phase-separating proteins, to further understand and generalize the guiding principles behind interfacial localization for distinct condensate targets. We repeated the optimization procedure for condensates based on the LAF-1 RGG domain and the N-terminal disordered domain of DDX4. The sequence compositions of the three targeted proteins are clearly distinct, as illustrated in Figure 5A, which shows the relative fraction of each amino acid category.^34^ Notably, hnRNPA1-LCD has the highest net charge among the three disordered proteins, with a net charge per residue (NCPR) of +0.057, although it contains relatively few charged residues and a higher proportion of aromatic residues compared to the other two cases. In contrast, LAF-1-RGG and DDX4N have fewer aromatic residues and lower net charges, with NCPR values of +0.024 and -0.017, respectively. The progression of the optimization targeting LAF-1-RGG was comparable to that of hnRNPA1-LCD, as measured by hypervolume improvement (Figure 5B). The majority of the gains occured in the first two iterations, after which the hypervolume reached a plateau. In contrast, the optimization for DDX4N was more challenging, as the Pareto front progressed more slowly and the hypervolume stagnated at a lower level.

**Figure 5:**
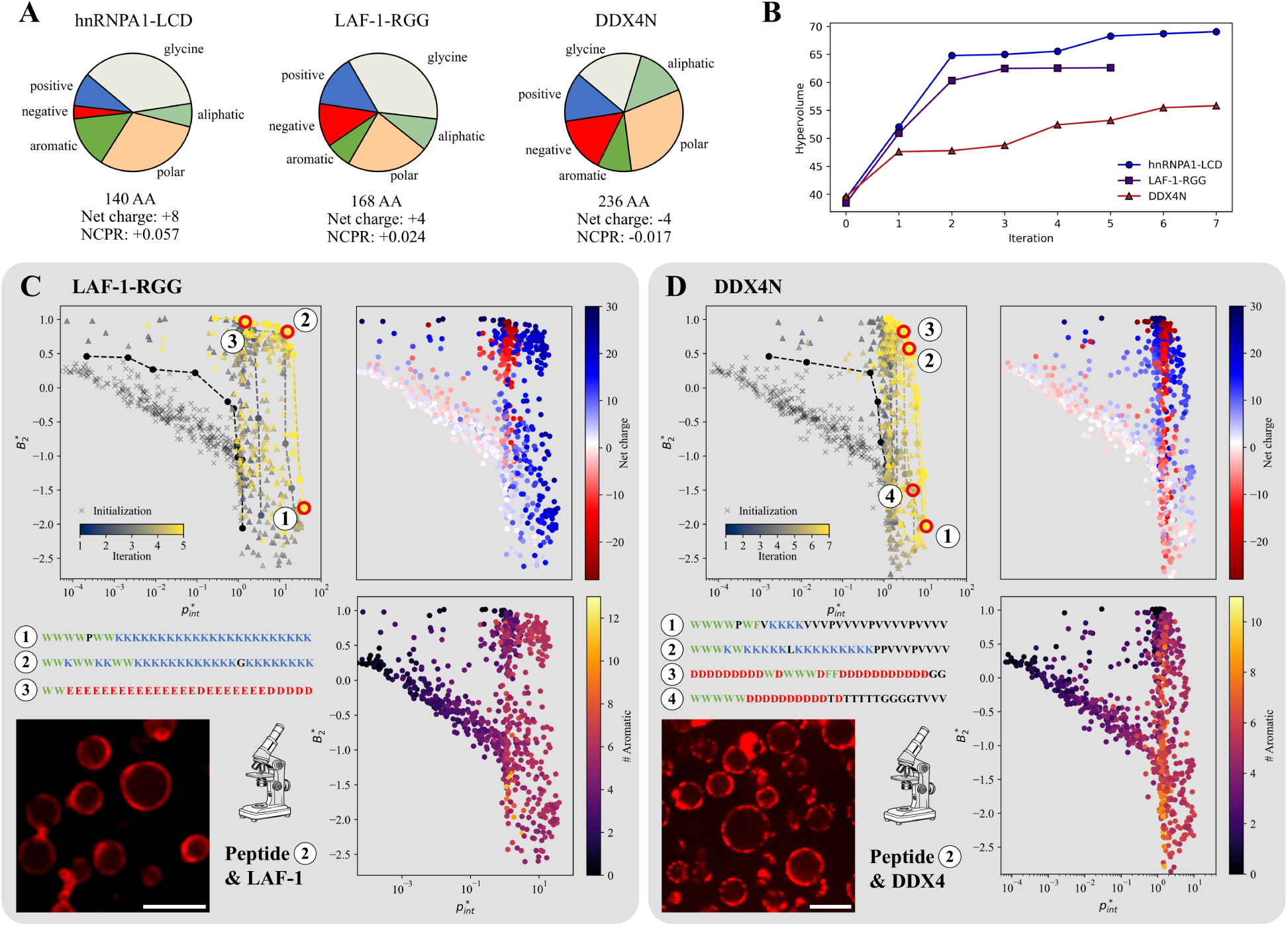
Expanding peptide design to other condensate targets: LAF-1-RGG and DDX4N. (A) Comparison of sequence composition among the three proteins of interest. hnRNPA1-LCD has the highest net charge per residue (NCPR) and the largest fraction of aromatics. In contrast, LAF-1-RGG and DDX4N both have fewer aromatic residues and lower net charges, with DDX4N showing the lowest NCPR. (B) Hypervolume progression for the peptide optimizations. (C) Peptide sequence optimization for LAF-1-RGG leads to surfactant-like peptides, mainly composed of tryptophans and lysines, where the overall composition remains nearly constant but the sequence patterning varies along the Pareto front. Interfacial localization of a selected peptide could be confirmed using confocal microscopy. (D) Peptide sequence optimization for DDX4N mostly yielded sequences featuring valine-rich tails and polylysine patches, resulting in a net charge opposite to that of the condensate. Experiments also confirmed interfacial localization *in vitro*. Scale bar: 5 *µ*m.

Similarly to the results obtained for hnRNPA1-LCD, the LAF-1-RGG peptide sequences exhibited a surfactant-like architecture (Figure 5C), with one end enriched in tryptophans and the other in lysines. The lysine-rich region was excluded from the condensate phase in the simulations. An important difference from the hnRNPA1-LCD case is that the peptides do not contain any arginine residues, potentially due to the low fraction of aromatic residues in LAF-1-RGG and the consequently lower relevance of cation–*π* interactions. Furthermore, the highest values for 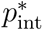 are lower, and the polylysine tail is longer than for hnRNPA1-LCD, which could be a consequence of the lower charge density of LAF-1-RGG condensates and thus a weaker driving force to exclude lysine residues. Another interesting observation is that the aromatic fraction and net charge remained almost constant along the main section of the Pareto front, covering a wide range of homotypic interaction strengths. The sequences mainly differed in patterning, as the aromatic patches become increasingly interrupted by self-repulsing lysines, which can be seen by for instance comparing peptide 1 and 2 in Figure 5C. Splitting up patches of strongly interacting “sticker” residues and therefore decreasing their valency is known to decrease interaction strength.^64^ Additionally, some sequences with high negative charge appeared in the Pareto front, exhibiting a slight preference for the interface over the dense phase (Table S3). This could be due to a combination of favorable long-range electrostatic interactions but insufficient short-range attraction, leading to the observed behavior. As previously done with the hnRNPA1-LCD condensates, we assessed the spatial localization of a selected peptide in LAF-1-RGG condensates using fluorescence confocal microscopy. The selected peptide also localized at the interface, as indicated by the fluorescent rim (Figure 5C). In contrast, a control peptide from the initialization did not partition at the interface and was instead homogeneously distributed within the condensate (Figure S6). The effect on the size distribution was similar to the hnRNPA-1-LCD case, with the condensates being smaller in presence of the peptide partitioning at the interface (Figure S8).

In the case of DDX4N (Figure 5D), the optimization results differed significantly from the other two cases. Most Pareto-optimal sequences did not exhibit a tail matching the condensate’s net charge, as observed in the previous cases, even though such architectures ere explored during the optimization process. One possible explanation for this behavior is the minimal net charge of DDX4N. Instead, the peptide sequences often featured valine-rich tails—which interact only weakly with most amino acids according to the Mpipi force field^28^—along with polylysine patches. While lysine exhibits favorable long-range electrostatic interactions with the condensate, its cation–*π* interactions are substantially weaker than those involving arginine, possibly aiding interfacial localization. This strategy appeared to perform better compared to variants matching the condensate’s net charge, such as peptide 4, which is not Pareto-optimal. Again, in agreement with *in silico* predictions, a peptide selected among the Pareto-optimal sequences did exhibit preferential interfacial partitioning *in vitro* (Figure 5D), while a control peptide spread uniformly in the condensate (Figure S6). As in the previous cases, interfacial partitioning also led to a decrease in DDX4N condensate size (Figure S8).

## Conclusions

In this study, we developed and applied a computational pipeline that integrates coarsegrained molecular simulations, machine learning, and mixed-integer linear programming (MILP) to de novo design peptides that partition at the interface of biomolecular condensates. The pipeline quantifies both the interfacial localization of isolated peptides and their homotypic interactions using coarse-grained simulations, followed by the training of a neural network and the identification of globally optimal sequences via bi-objective MILP. This approach led to sets of self-repulsive peptides with a high propensity to partition at the interface of three distinct condensates based on hnRNPA1-LCD, LAF-1-RGG, and DDX4N. Interfacial accumulation of the designed peptides was experimentally confirmed *in vitro* for all three cases, validating the design approach. To our knowledge, this is the first study showing that short (30-residue long) peptides can preferentially partition at the interface of the condensates.

The peptide sequences generally exhibited surfactant-like architectures with two tails, one interacting with the scaffold molecule of the condensate and a second one excluded from the dense phase (Figure 6). This architecture is similar to the amphiphilic nature of much larger proteins previously shown to partition at the interface of biomolecular condensates.^7,10–12,22–25^ In all cases, the tail of the peptide entering the dense phase was enriched in aromatic residues (and also in arginine for the hnRNPA1-LCD case), to promote attractive interactions with the scaffold molecule of the condensates. In contrast, the tail that was excluded from the condensates was different for the distinct IDRs. In the case of hnRNPA1-LCD and LAF-1-RGG, the tail primarily comprised lysine residues and matched the net charge of the condensate-forming protein, therefore promoting electrostatic repulsive interactions with the scaffold molecule of the condensates. This polylysine tail was longer for LAF-1-RGG, likely due to the lower positive charge density of the protein. In the case of DDX4N, which has the lowest charge density of the three protein targets, the mechanism behind the exclusion of the tail from the condensates cannot be based on electrostatic repulsion. Instead, interactions with the scaffold molecules were minimized by introducing in the excluded tail uncharged valine residues, which exhibit negligible interactions with the scaffold protein. These findings indicate that the net charge of the condensate-forming protein is a key physicochemical feature for designing peptides exhibiting specific interfacial localization.

**Figure 6:**
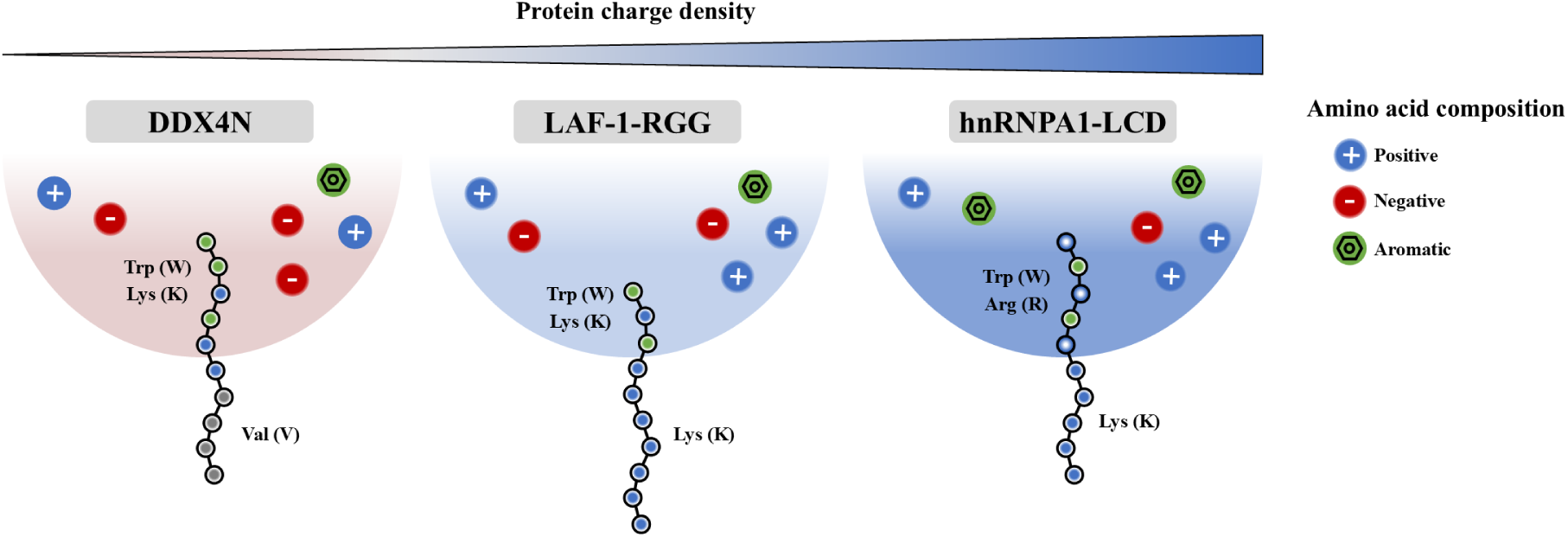
Schematic illustration of the design outcome for interface-partitioning peptides based on distinct condensate targets. The resulting surfactant-like peptide sequences indicate the charge density of the condensate-forming protein as a key physicochemical feature.

Overall, these findings and the developed pipeline provide a promising strategy for the rational design of interface-localizing peptides or protein sequences tailored to a specific condensate. The active learning framework using neural networks and MILP for sequence optimization can be applied to other design targets, avoiding getting trapped in local optima. In the future, the approach could be expanded to more complex condensate compositions, also involving globular domains and client molecules.

## Materials and Methods

### Molecular simulations

All molecular simulations were carried out using the Mpipi force field^28^ in an NVT ensemble applying a Langevin thermostat^65^ at 300 K with a relaxation time of 100 ps. Simulations were performed using the LAMMPS Molecular Dynamics package (version 2nd of August 2023)^66^ and the adaptive biasing force method was implemented using the Colvars module. ^67^

### Interface partitioning

The condensate dense phase was modeled using 16 protein copies within a periodic box, with dimensions 71×71×343 Å^3^ (hnRNPA1-LCD), 76×76×450 Å^3^ (LAF-1-RGG) and 88×88×510 Å^3^ (DDX4N), originating from a 2×2×4 starting configuration with density 0.5 g/cm^3^. The dense phase was loosely restrained in z direction using harmonic walls (force constant: 5 kcal mol^−1 °^A^−2^) to a maximum width of 145 ^°^A(hnRNPA1), 160 ^°^A(LAF-1) and 170 ^°^A(DDX4) preventing strong fluctuations of the interface or dissociation of proteins into the dilute phase. The small system size and harmonic walls enable faster simulations with acceptable statistical error. The z-component of the center-of-mass distance between the peptide and dense phase (hnRNPA1: 0-12 nm, LAF-1/DDX4: 0-12.8 nm) served as the collective variable for the adaptive biasing force method (ABF). This range was divided into four sections, resulting in four separate simulations whose PMFs were later combined. Two PMFs were generated per simulation, with one peptide positioned on each side of the slab, ensuring the peptides remained in separate sections where interactions are avoided. This process was replicated three times leading to a total of 6 PMFs obtained from 12 independent simulations for every peptide. The PMF was constructed using a total number of 64 bins, applying the biasing force after collecting 5,000 samples per bin. Each simulation began with energy minimization and 5 ns of equilibration, followed by a 250 ns production run. Δ*G*_1_ was determined as the mean difference between the PMF value at 0 nm (dense phase) and the global minimum, while Δ*G*_2_ was calculated as the free energy difference between the values at 0 nm (dense phase) and 12 nm (dilute phase).

### Homotypic interactions

Homotypic interactions were quantified by constructing the PMF as a function of the centerof-mass distance between two peptide copies, ranging from 0 to 70 ^°^A, using the adaptive biasing force method in a 140×140×140 Å^3^ simulation box with periodic boundary conditions. The bin width was set to 0.5 ^°^Aand the biasing force applied after collecting 500 samples per bin. Energy minimization and 1 ns of equilibration were followed by 1 *µ*s production simulation, repeated eight times for each peptide. The PMF was converted into the second virial coefficient *B*_2_ according to:^40,51^

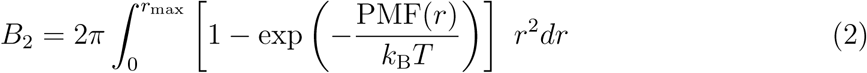

A bi-symmetric log transformation^52^ was applied to the average *B*_2_ for every peptide, preventing a highly skewed distribution of data and the associated difficulties with training a machine learning model (C = 20 nm^3^).

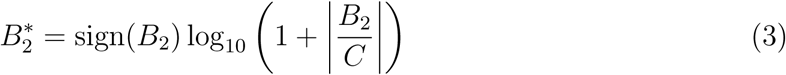

### Contact map simulation

The simulation setup for generating the contact map in Figure 3C was identical to the interface partitioning simulations for hnRNPA1, except without harmonic walls and adaptive biasing force. Energy minimization and 30 ns of equilibration were followed by 300 ns production simulation with coordinates saved every 50 ps. The contacts were anaylyzed using PLUMED^68^ (version 2.8.0) with a cutoff of 1.0 nm.

### Machine learning and optimization

After each round of simulations, a multi-output neural network with two fully connected hidden layers of 50 neurons each was trained on Δ*G*_1_, Δ*G*_2_, and *B*^∗^_2_ with PyTorch^69^ (version 2.3.1) and scikit-learn^70^ (version 1.5.1) using a set of engineered sequence features (Supplementary Information). All features and objectives were normalized to the range [-1, 1], and the Smooth L1 loss function was minimized with the Adam optimizer.^71^ A multi-step learning rate scheduler was employed, decaying the learning rate by 0.9 at epochs 200, 400, 600, and 800. A batch size of 32 was used for training. 10% of the training data was set aside as a validation set, and the model exhibiting the lowest validation loss was selected. In every iteration, an 80/20 train–test split was used to optimize hyperparameters (initial learning rate and weight decay). The performance of the best model was then evaluated on a separate holdout set (20%). The final model used for optimization was trained on the entire available dataset, after which pruning was applied: all weights and biases of the neural network with absolute values below 0.001 were set to zero, mitigating potential numerical issues when solving the MILP. A MILP formulation of the trained neural network was incorporated into the algebraic modeling framework Pyomo^56,57^ (version 6.7.3) using OMLT^44^ (version 1.1). In the first step, the two objectives, ln(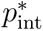) and *B*^∗^_2_, were separately optimized with Gurobi,^58^ using the one-hot encoded amino acid sequence as the input. Details on integrating the AGGRESCAN^47^ predictor as a constraint and on converting Δ*G*_1_ and Δ*G*_2_ into ln(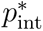) are provided in the Supplementary Information. After determining the bounds, the *ε*-constrained method was applied by constraining *B*^∗^_2_, resulting in 50 Pareto-optimal peptide sequences (exploitation). 25 well-spaced points were selected for simulation in the next round. For the exploration part, the weighted-sum method described in the main text was used to generate 150 sequences, from which a well-spaced subset of 75 sequences was selected. The weighted-sum method defines a combined objective using random weights assigned to the individual objectives, which differ for each exploration point. Furthermore, between 2 and 20 positions (chosen uniformly at random) were constrained to randomly selected amino acids. The remaining positions were then optimized. This approach strikes a balance between exploration and sampling relevant regions of the design space. In the first optimization with hnRNPA1-LCD as the condensate target, we reduced the number of exploration points to 25 for iterations 3 and 4, and to zero for the last three iterations, fully focusing on the exploitation component.

### Protein and peptide production

Genes encoding the different LCDs were codon optimized for expression in E. coli, synthesized and cloned between NdeI and BamHI restriction sites into the pET-15b vector by Genewiz (NJ, US). Recombinant proteins were produced in E. coli BL21(DE3) GOLD cells. Bacterial cultures were induced at OD 0.7 with 0.5 mM isopropyl D-thiogalactopyranoside and grown for an additional 16 h at 20 °C (DDX4N) or 37 °C (LAF-1-RGG and hnRNPA1-LCD). Cells expressing the proteins were centrifuged, re-suspended in lysis buffer (LAF-1-RGG/hnRNPA1-LCD: 8 M urea, 1 M NaCl, 10 mM imidazole, 50 mM Tris at pH 7.5, 2 mM *β*-mercaptoethanol; DDX4N: 1 M NaCl, 10 mM imidazole, 50 mM Tris at pH 8.5, 2 mM *β*-mercaptoethanol), lysed by sonication, and centrifuged at 18,300 g for 20 min. The supernatant was further used for purification of the proteins using immobilized-metal affinity chromatography (elution buffers: 1 M NaCl, 500 mM imidazole, 2 M urea; 50 mM Tris at pH 7.5, 2 mM *β*-mercaptoethanol for LAF-1-RGG/hnRNPA1-LCD and 1 M NaCl, 500 mM imidazole, 50 mM Tris at pH 8.5, 2 mM *β*-mercaptoethanol for DDX4N). Size exclusion chromatography using a Superdex 75 16/600 column (GE Healthcare) in 50 mM Tris at pH 7.5, 500 mM NaCl, 2M urea, 10% glycerol for hnRNPA1-LCD; 50 mM Tris at pH 7.5, 500 mM NaCl, 2M urea for LAF-1-RGG and 50 mM Tris at pH 8.5, 1 M NaCl, 10% glycerol for DDX4N was used as the final purification step. The quality of the purified proteins was checked by SDS-PAGE electrophoresis. Peptides were purchased from GenScript with a purity ≥ 95%. The Cy5 dye was conjugated to the N-terminus, while the C-terminus was amidated.

### Brightfield and confocal microscopy

The different LCDs (10 *µM* ) were incubated with either the corresponding interface peptide or a control peptide at a protein-to-peptide ratio of 10:1. For hnRNPA1-LCD, the buffer composition was 20 mM Tris, 150 mM NaCl, pH 7.5, whereas for LAF-1-RGG and DDX4N, the buffer consisted of 20 mM Tris, 25 mM NaCl, pH 7.5. Interface partitioning was assessed by measuring the intrinsic fluorescence of Cy5, excited with a 633 nm laser, with emission recorded at 670–700 nm, using a confocal microscope (Leica TCS SP8) equipped with a 63× NA 1.4 oil objective (Leica). Imaging was performed with the Leica Application Suite X (LAS X) software, version 1.0. The fluorescence intensity profiles across the condensates were estimated using ImageJ.^72^

Brightfield microscopy was used to estimate the size of protein condensates in the absence and presence of the different interface peptides. The different LCDs were incubated at 10 *µM* , with and without the corresponding interface peptides, at a protein-to-peptide ratio 10:1 in the corresponding buffer. After sedimentation, the condensates were visualized using a brightfield microscope (Eclipse Ti-E, Nikon) using a 60x oil objective (FI Plan Apo Lambda NA 1.4, Nikon). Their size distribution was analyzed using ImageJ. All samples were analyzed in 384-well plates (MatriPlate 384-Well Plate, Glass Bottom, Brooks).

### Dynamic light scattering (DLS)

The size distribution of hnRNPA1-LCD condensates in the dilute phase, both in the absence and presence of peptide, was estimated using a Zetasizer Nano-ZS (Malvern) at 25°C. hnRNPA1-LCD at 10 *µM* was incubated with and without the interface peptide at a proteinto-peptide ratio of 10:1 in 20 mM Tris, 150 mM NaCl, pH 7.5. After 1 h of incubation, the samples were centrifuged at 8,000 × *g* for 30 minutes to separate the dilute and dense phases.

## Data availability

Pareto fronts for all design cases, ML benchmarks and final model performances, benchmarks with GA, as well as microscopy and DLS data are provided in the Supplementary Information.

## Acknowledgements

We kindly acknowledge the European Research Council through the Horizon 2020 research and innovation programme (grant agreement No. 101002094) for financial support. Computations were performed on the high-performance computing cluster Euler by ETH Zurich. We acknowledge Florence Stoffel for assistance with the experiments.

## Author contributions

T.N.S. and P.A. designed the conceptual framework of the study. T.N.S. and M.A.B. developed and applied the computational pipeline. M.G. and L.F. performed the experiments. L.F.S. and G.G. supervised the machine learning and MILP parts. T.N.S, M.G., M.A.B., and P.A. wrote the manuscript with contributions from all authors. P.A. acquired funding and supervised the project.

