## Supplementary Information for "De novo design of peptides localizing at the interface of biomolecular condensates"

### 1 Machine learning models

#### 1.1 Featurization of peptide sequences

Given the relatively limited amount of data (300+ sequences) compared to the massive design space, utilizing engineered descriptors has proven to be an effective approach to represent polymer or protein sequences.[1,2] For the optimization to work with an MILP solver, the featurization has to be a linear mapping of the amino acid sequence (30x20 one-hot matrix) to the feature representation. 44 handcrafted features were used to represent the amino acid sequence in a numerical fashion. These features were constructed as follows:

- 20 features are designated to represent the composition by quantifying the count of amino acids of type X within the sequence.

- To represent positive and negative charge patterning we utilize zeroth to third moment of the positive and negative charge distribution along the sequence.

$$P_n = \sum_{i=1}^{N} \text{charge}_i^+ * d_i^n \qquad n \in [0, 3] \tag{1}$$

$$N_n = \sum_{i=1}^{N} \text{charge}_i^- * d_i^n \qquad n \in [0, 3] \tag{2}$$

  where $d$ represents the positional distance of residue $i$ from the center of the amino acid sequence, $d_i \in [-14.5, 14.5]$ for the case of a 30 amino acid long peptide. The zeroth moments ($n = 0$) represent the total number of positive and negative charges, respectively. The same concept is used for the distribution of aromatic residues and molecular weight:

$$A_n = \sum_{i=1}^{N} \text{aromatic}_i * d_i^n \qquad n \in [0, 3] \tag{3}$$

$$MW_n = \sum_{i=1}^{N} MW_i * d_i^n \qquad n \in [0, 3] \tag{4}$$

where aromatic$_i$ is a binary variable we define as one for phenylalanine, tryptophan and tyrosine and as zero for all other residues. We also assign the parameter $\epsilon$ to each residue quantifying homotypic van der Waals interactions, inspired from the original Mpipi paper:[3]

$$\epsilon = \int_{\sigma}^{3\sigma} \phi(r)dr \tag{5}$$

where $\phi(r)$ is the Wang-Frenkel potential[4] with residue-specific parameters. We then use $\epsilon$ to define four more features:

$$E_n = \sum_{i=1}^{N} \epsilon_i * d_i^n \qquad n \in [0, 3] \tag{6}$$

- We also include an adapted sequence hydropathy decoration, SHD, which was adapted from the original SHD[5] due to the absence of a hydropathy value $\lambda$ in the Mpipi force field:

$$SHD = \sum_{i=1}^{N} \sum_{j=i+1}^{N} (\epsilon_i + \epsilon_j)(j - i)^{-1} \tag{7}$$

- The final features originate from the sequence charge decoration SCD.[6] Due to the non-linear nature of the original version, we defined three similar features that can be utilized in a MILP formulation:

$$SPD = \sum_{i=1}^{N} \sum_{j=i+1}^{N} (\text{charge}_i^+ + \text{charge}_j^+)(j - i)^{-1} \tag{8}$$

$$SND = \sum_{i=1}^{N} \sum_{j=i+1}^{N} (\text{charge}_i^- + \text{charge}_j^-)(j - i)^{-1} \tag{9}$$

$$SPND = \sum_{i=1}^{N} \sum_{j \neq i}^{N} (\text{charge}_i^+ + \text{charge}_j^-)|j - i|^{-1} \tag{10}$$

It is important to note that the results of the simulations are invariant to sequence flipping due to the coarse-grained representation. To account for this invariance, a preprocessing step was implemented: if the third moment of the molecular weight ($MW_3$) was negative, the sequence was flipped before computing the descriptor set. In the MILP optimization, a constraint $MW_3 \geq 0$ was added to enforce this condition. This procedure effectively reduced the design space to $\sim 20^{30}/2$. In the final Pareto front, the sequences were again flipped to ensure that the N-terminal aromatic Cy5 dye was attached to the tail prone to condensate interactions.

#### 1.2 Benchmarking machine learning models

Using the initial data from the hnRNPA1-LCD optimization, we evaluated the performance of a fully connected two-layer multi-output neural network with varying layer widths and compared it against alternative models, including elastic net (EN), support vector machines (SVM), and gradient-boosted trees (GBT) (Figure S1). The predictive performance was evaluated by computing the coefficient of determination ($R^2$) on 20 randomly generated train-test splits, using an 80/20 split ratio. The neural network training procedure followed the methods described in the main text, with the only difference being the use of fixed values for the initial learning rate ($10^{-3}$) and weight decay ($10^{-3}$). A grid search was performed to tune hyperparameters for the SVM and EN models (EN: L1 ratio; SVM: $C$, $\epsilon$), with each combination evaluated using 5-fold cross-validation. For the gradient boosting regressor, a randomized search over 200 models was conducted to optimize hyperparameters (learning rate, number of estimators, loss function, splitting criterion, minimum samples for splitting, maximum tree depth, and minimum samples per leaf), also assessed through 5-fold cross-validation. All models were implemented using scikit-learn[7] and PyTorch.[8] We observed that the neural network's performance plateaued above 40 nodes per layer and that it generally outperformed elastic net, support vector machines, and gradient-boosted trees. To maintain some flexibility for future data, we chose a layer width of 50 to go forward.

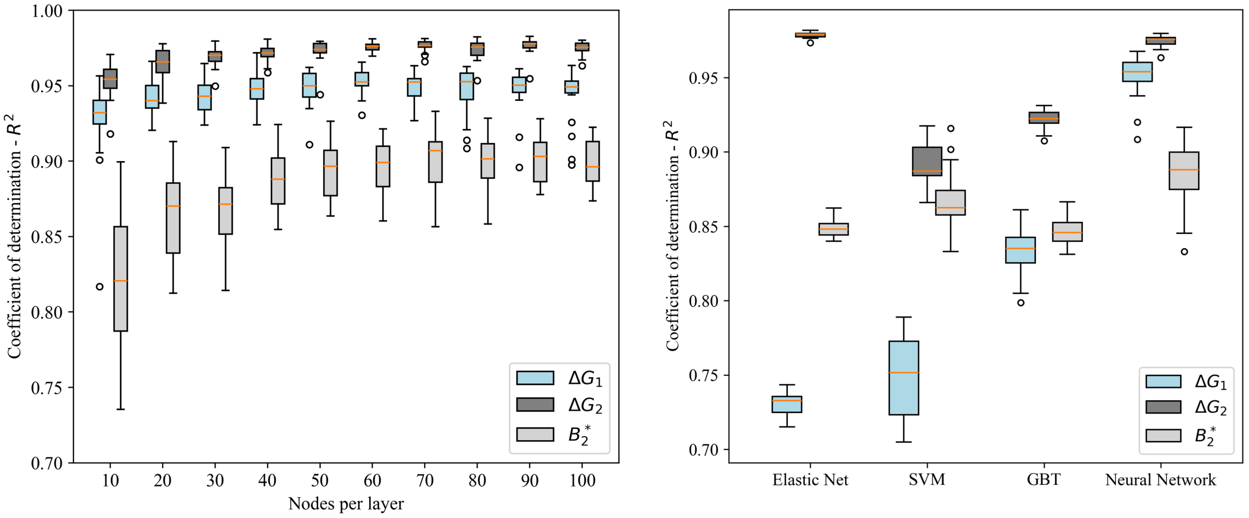


Fig. S1: Benchmarking the performance of a two-layer neural network with varying layer widths against alternative models (elastic net, support vector machines, and gradient-boosted trees), using the initialization data for the hnRNPA1-LCD condensate target. Based on these results, a neural network width of 50 was chosen.

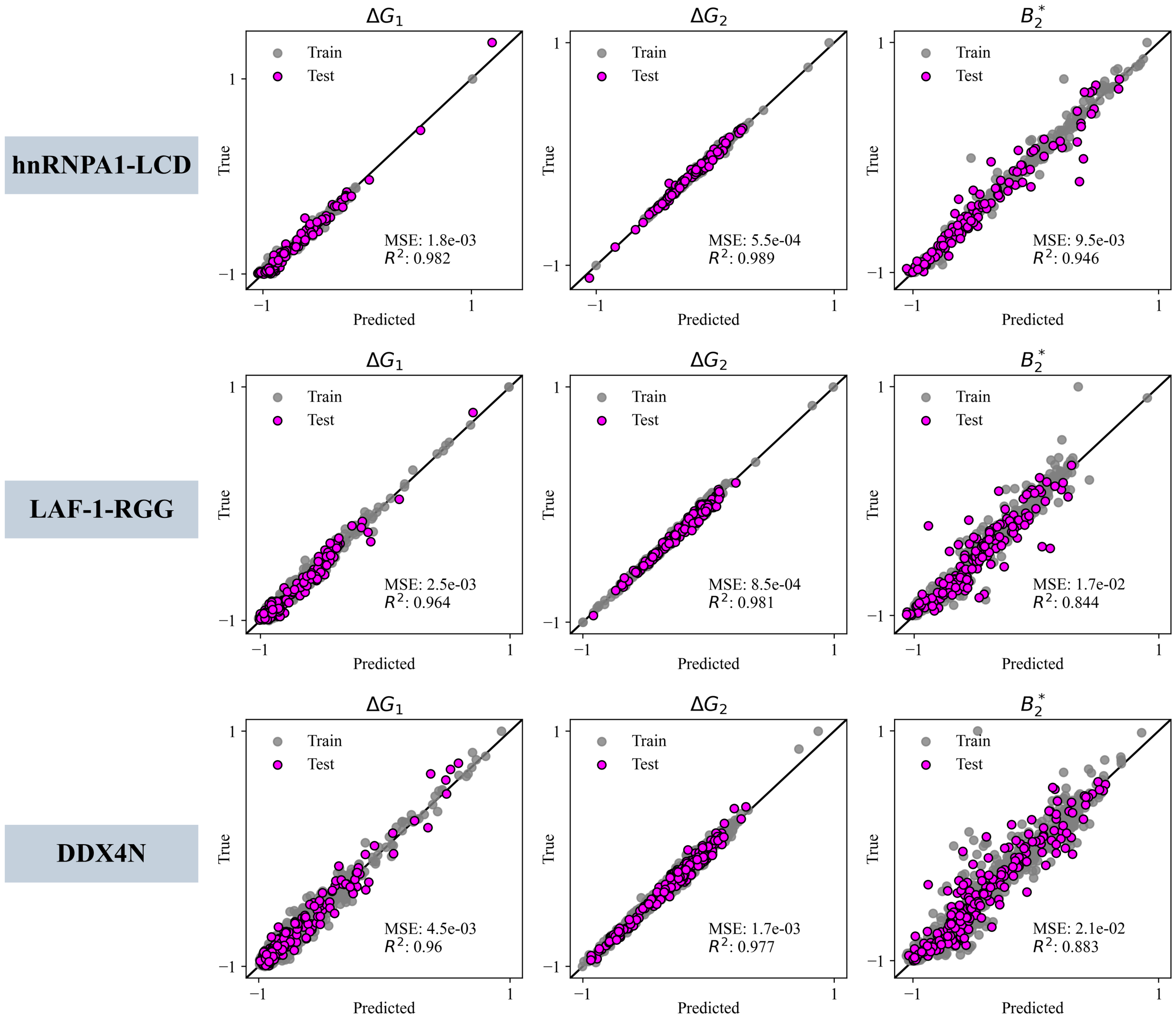


Fig. S2: Performance of the final model on a 20% test set, evaluated after completing all iterations in each optimization.

### 2 Objective function interface partitioning

We defined the objective of maximizing interface partitioning as maximizing the ratio of the peptide's probability of localizing at the interface $p_{\text{int}}$ to the probabilities of localizing in dense $p_{\text{den}}$ or dilute phase $p_{\text{dil}}$:

$$\max\left[\frac{p_{\text{int}}}{p_{\text{den}}+p_{\text{dil}}}\right] = \max\left[\frac{p_{\text{den}}}{p_{\text{int}}}+\frac{p_{\text{dil}}}{p_{\text{int}}}\right]^{-1} \tag{11}$$

which can be reformulated using the potential of mean force $W$ as

$$\max\left[\frac{\int_{\text{den}}\exp\left(-\frac{W(q)}{k_{\text{B}}T}\right)dq}{\int_{\text{int}}\exp\left(-\frac{W(q)}{k_{\text{B}}T}\right)dq}+\frac{\int_{\text{dil}}\exp\left(-\frac{W(q)}{k_{\text{B}}T}\right)dq}{\int_{\text{int}}\exp\left(-\frac{W(q)}{k_{\text{B}}T}\right)dq}\right]^{-1} \tag{12}$$

When approximating constant $W$ for dilute and dense phase and interface, and that $W_{\text{int}}$ is given by the minimum in the PMF, we can simplify to

$$\begin{aligned}\max\left[\frac{V_{\text{den}}}{V_{\text{int}}}\exp\left(-\frac{W_{\text{den}}-W_{\text{int}}}{k_{\text{B}}T}\right) + \frac{V_{\text{dil}}}{V_{\text{int}}}\exp\left(-\frac{W_{\text{dil}}-W_{\text{int}}}{k_{\text{B}}T}\right)\right]^{-1} \equiv \\ \equiv \max\left[\exp\left(-\frac{\Delta G_1}{k_{\text{B}}T}\right) + \phi\exp\left(-\frac{\Delta G_1+\Delta G_2}{k_{\text{B}}T}\right)\right]^{-1}\end{aligned} \tag{13}$$

where $\phi$ denotes the volume ratio of the dilute to dense phases, which we approximate as 100. This corresponds to a rough estimate and is system dependent, and based on initialization results suggesting that tuning $\Delta G_2$ is considerably easier than tuning $\Delta G_1$, we intentionally overestimated the second term, giving rise to our final objective to be maximized:

$$p^*_{\text{int}} = \left[\exp\left(-\frac{\Delta G_1}{k_{\text{B}}T}\right) + \phi\exp\left(-\frac{\Delta G_2}{k_{\text{B}}T}\right)\right]^{-1} \tag{14}$$

In order to reduce skewedness and associated numerical issues, we minimized $\ln(1/p^*_{\text{int}})$, which is an equivalent optimization problem to maximizing $p^*_{\text{int}}$. $\ln(1/p^*_{\text{int}})$ is also still a convex function with respect to the variables $\Delta G_1$ and $\Delta G_2$, which can be demonstrated by applying Sylvester's criterion.[9] In other words, let the objective function $f : \mathbb{R}^2 \rightarrow \mathbb{R}$ with variables $x$ and $y$ be

$$f(x,y) = \ln\left(\exp(-x) + \phi\exp(-y)\right) \quad x, y \in \mathbb{R},\ \phi \in \mathbb{R}_{>0}$$

We can then calculate the Hessian matrix ($H$), all principal minors ($a_{11}$ and $a_{22}$), and the determinant ($\det(H)$):

$$\begin{aligned}
H &= \begin{bmatrix} \frac{\exp(-x)}{\exp(-x)+\phi\exp(-y)} - \frac{\exp(-2x)}{(\exp(-x)+\phi\exp(-y))^2} & -\frac{\phi\exp(-x-y)}{(\exp(-x)+\phi\exp(-y))^2} \\ -\frac{\phi\exp(-x-y)}{(\exp(-x)+\phi\exp(-y))^2} & \frac{\phi\exp(-y)}{\exp(-x)+\phi\exp(-y)} - \frac{\phi^2\exp(-2y)}{(\exp(-x)+\phi\exp(-y))^2} \end{bmatrix} \\
&= \begin{bmatrix} a_{11} & a_{12} \\ a_{21} & a_{22} \end{bmatrix} \\
a_{11} \geq 0 &\iff \frac{\exp(-x)}{\exp(-x)+\phi\exp(-y)} \geq \frac{\exp(-2x)}{(\exp(-x)+\phi\exp(-y))^2} \\
&\iff \exp(-y) \geq 0 \quad \forall\{x,y\} \in \mathbb{R} \\
a_{22} \geq 0 &\iff \exp(-x) \geq 0 \quad \forall\{x,y\} \in \mathbb{R} \\
\det(H) &= a_{11}a_{22} - a_{12}a_{21} = 0 \quad \forall\{x,y\} \in \mathbb{R}
\end{aligned}$$

Because all principal minors and the determinant of the Hessian are non-negative, the matrix is positive semi-definite, and therefore the objective function $f$ is convex. This convexity allowed to approximate the objective function by adding linear first order Taylor approximations to the MILP, the principle is illustrated in Figure S3 for a univariate function.[10] The linearization was performed in a gridwise fashion for 30 points in both $\Delta G_1$ and $\Delta G_2$ ($S_1$ and $S_2$, respectively), ranging from the smallest value to the largest value obtained in the training data, resulting in a total of 900 supporting planes. For a single point $s_1 \in S_1$ and $s_2 \in S_2$, we introduced the following constraint for the linearized interface partitioning objective ($\ell \approx \ln(1/p^*_{\text{int}})$) to be minimized:

$$\begin{aligned}\ell \geq \ln\left[\exp\left(-\frac{s_1}{k_\text{B}T}\right) \;+\; \phi\exp\left(-\frac{s_2}{k_\text{B}T}\right)\right] \\ -\; \frac{\exp\left(-\frac{s_1}{k_\text{B}T}\right)}{\exp\left(-\frac{s_1}{k_\text{B}T}\right) \;+\; \phi\exp\left(-\frac{s_2}{k_\text{B}T}\right)}\frac{1}{k_\text{B}T}\left(\Delta G_1 \;-\; s_1\right) \\ -\; \frac{\phi\exp\left(-\frac{s_2}{k_\text{B}T}\right)}{\exp\left(-\frac{s_1}{k_\text{B}T}\right) \;+\; \phi\exp\left(-\frac{s_2}{k_\text{B}T}\right)}\frac{1}{k_\text{B}T}\left(\Delta G_2 \;-\; s_2\right)\end{aligned} \tag{15}$$

By adding this constraint for all combinations of $s_1$ and $s_2$, the nonlinear objective (14) was reformulated to be compatible with mixed-integer linear programming, such that:

$$\max\; p^*_{\text{int}} \;\approx\; \begin{array}{ll} \min & \ell \\ \text{s.t.} & \text{Eq (15)}, \quad \forall s_1 \in S_1, s_2 \in S_2 \end{array} \tag{16}$$

Notice that, minimizing $\ell$ combined with the direction of the inequality ($\geq$) in (15) favors the constraint to be active and, therefore, enforces the linear approximation without the need for equality constraints.

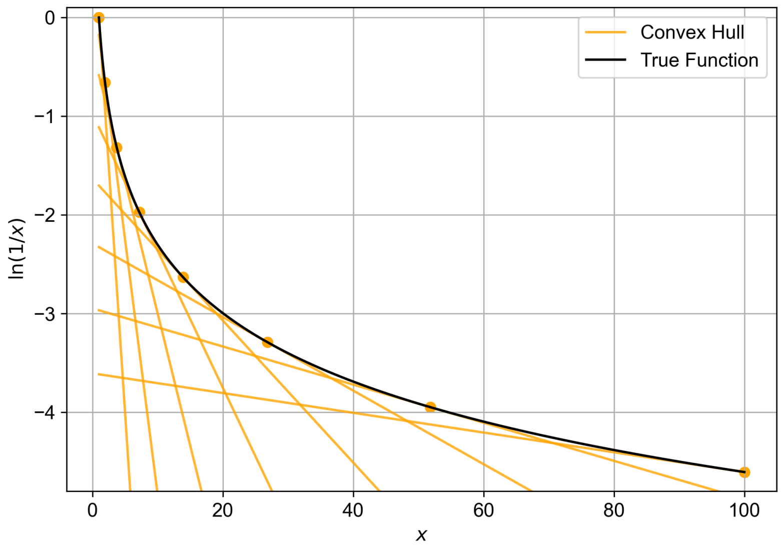


Fig. S3: Illustration of the convex hull approach.

### 3 Integration of AGGRESCAN predictor

We incorporated AGGRESCAN as a constraint in the MILP optimization because its prediction algorithm is based on linear operations, making it suitable for integration into an MILP. For all details about the algorithm, we refer to the original publication.[11] AGGRESCAN calculations are based on aggregation-propensity values per amino acid (aaAV, or a3v). The a3v is averaged with a sliding window of length 5 given the sequence length of 30, resulting in an a4v value assigned to the central residue in each window. If there are 5 or more sequential residues with an a4v larger than the hot spot threshold (HST$= -0.02$) and none of the amino acids inside this window is a proline, the sequence contains an aggregation hot spot and is thus defined as infeasible in the MILP. To formulate this, one has to find a suitable linearization for finding the minimum of a list of 5 values, which should not exceed the HST. This was achieved by introducing big-M constraints. We consider the following equation:

$$X = \min\{x_1, ..., x_n\} \tag{17}$$

For this we have to introduce $n$ binary variables $z_i$ which are equal to 0 if the value at position $i$ is the minimum and 1 otherwise. We also introduce a parameter $M$ which has to be larger than the largest value inside the list, but as small as possible. We can then introduce the following constraints:

$$X \geq x_i - M * z_i \qquad i \in n \tag{18}$$

$$X \leq x_i \qquad i \in n \tag{19}$$

$$\sum_{i}^{n} z_i = n - 1 \tag{20}$$

These constraints were implemented for each window combined with $X \leq$ HST, ensuring that the minimal value obtained was less than or equal to the HST. If a window contained a proline, we ensured that $X$ was lower than the HST by subtracting 7, always leading to $X <$ HST.

### 4 Comparing MILP and genetic algorithm

Mixed-integer linear programming (MILP) can generate the true Pareto front, whereas genetic algorithms (GA) risk becoming trapped in local optima. To investigate this in our optimization case, we constructed the Pareto front using trained neural networks for the first two iterations with hnRNPA1-LCD as a condensate target. As a genetic algorithm, we employed NSGA-II,[12] utilizing genetic operations at the sequence level (Figure S4):

- Point mutations randomly replace a single amino acid with any other amino acid, each with equal probability.
- Crossover events exchange subsequences between sequences at a randomly chosen split position.

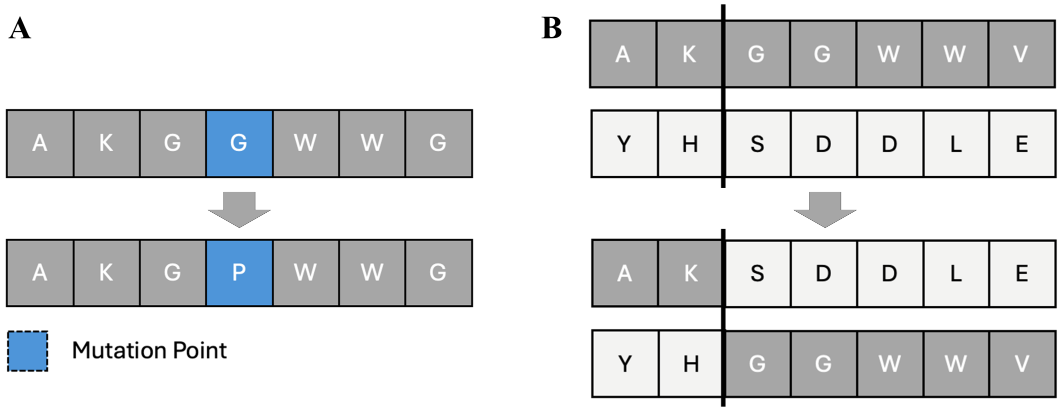


Fig. S4: Genetic operations employed in the GA were point mutations (A) and crossover events (B).

NSGA-II was implemented using a generation size of 200. The probabilities for both point mutation and crossover event were set to 50%. As an initial population, the 300 random sequences from the initialization were used. The AGGRESCAN predictor was implemented as a constraint in the following manner: if a genetic operation lead to a constraint violation, the sequence was discarded and a new sequence is generated. This ensured that at no point any aggregation-prone sequence was produced. The genetic algorithm was run until the hypervolume of the Pareto front stagnated.

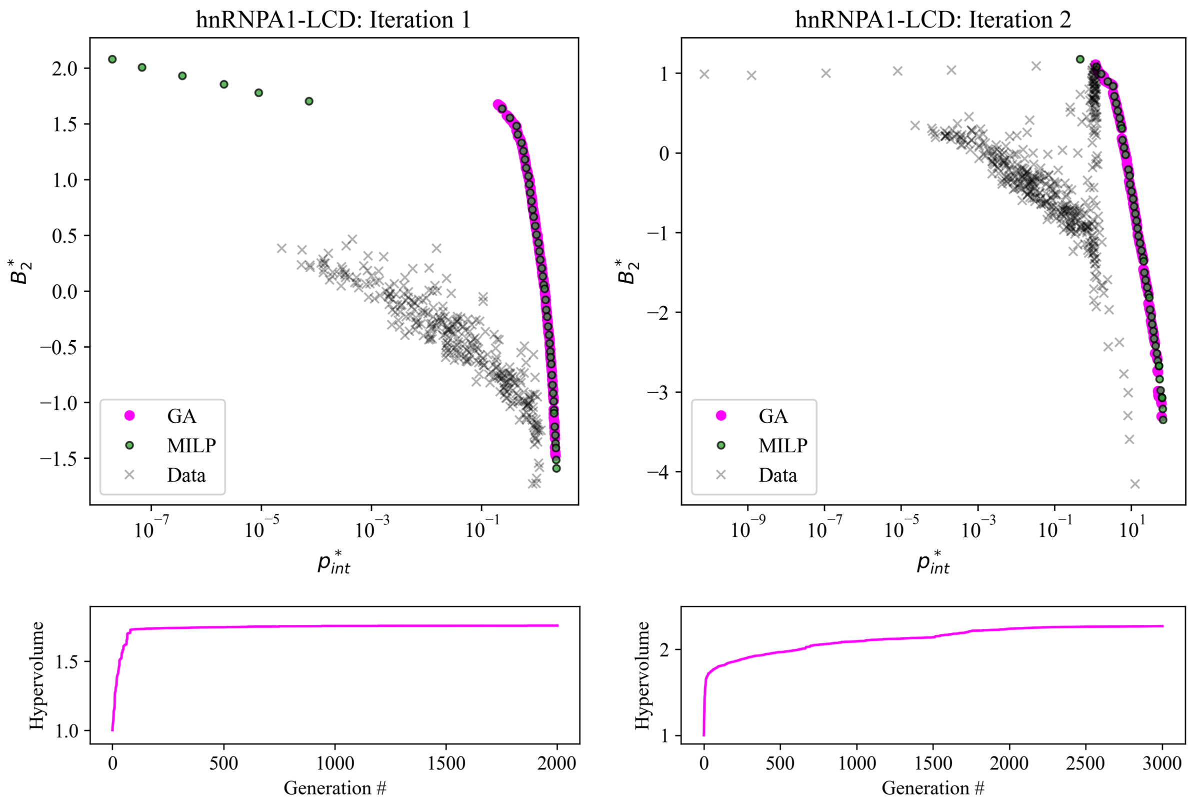


Fig. S5: Comparison of the genetic algorithm (GA) and mixed-integer linear programming (MILP) in constructing Pareto fronts of the first two trained surrogate models in the hnRNPA1 optimization case. While the GA captured most of the true Pareto front identified by MILP for both models, it failed to recover a significant portion in the iteration 1 model, despite hypervolume convergence.

To compare the capability of GA and MILP in generating Pareto fronts, we applied both methods to two trained surrogate models: one based on the initialization data (used for iteration 1) and another after one additional iteration. It is important to note that no actual iterations were performed using the GA; only the surrogate models at these stages were optimized. While the genetic algorithm reconstructed most of the true Pareto front identified by MILP, it failed to capture a significant portion with high $B_2^*$ values in the iteration 1 model (Figure S5). Although the lack of hypervolume improvement with increasing generations suggested convergence, the GA was in fact trapped in a local optimum. This highlights the superior performance of MILP for this optimization problem.

### 5 Confocal microscopy

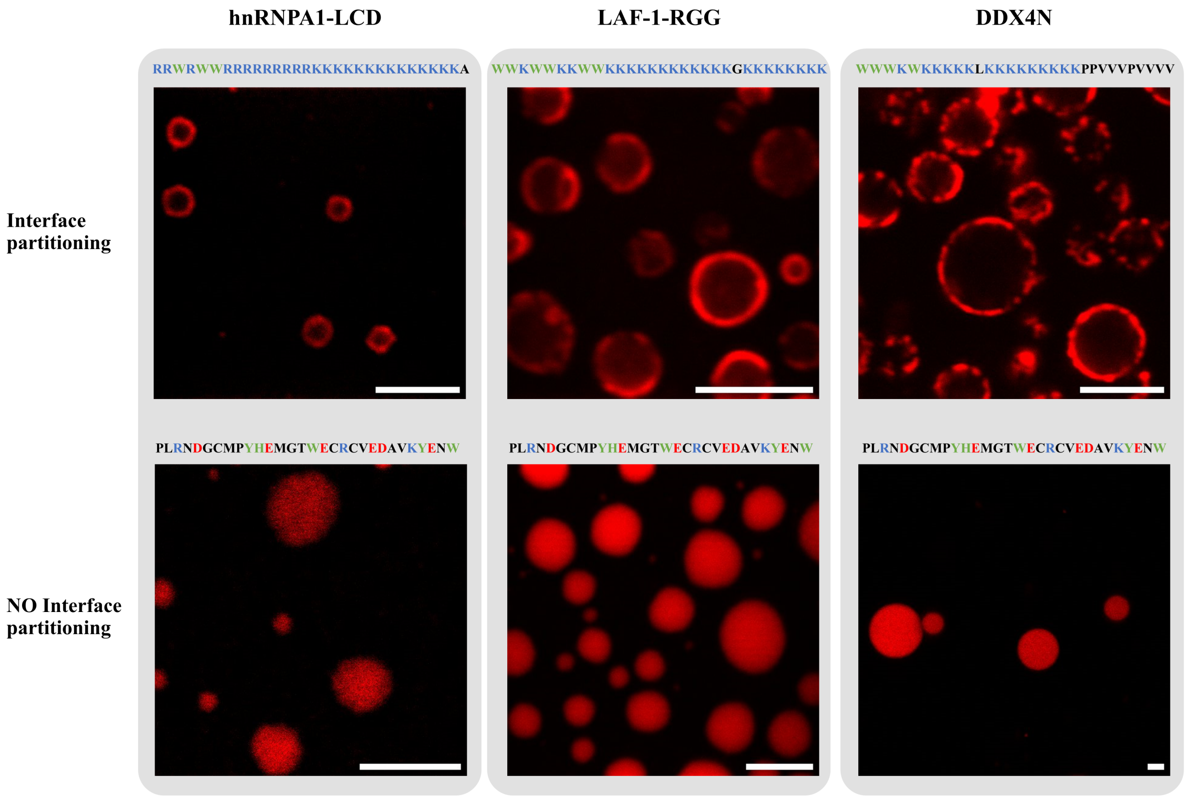


Fig. S6: Experimental validation of interface partitioning at condensates formed by the proteins hnRNPA1-LCD, LAF-1-RGG and DDX4N. Fluorescence confocal microscopy confirmed that peptides with high predicted preference for the interface accumulated at the condensate interface (top), while the control peptide spread uniformly (bottom). Scale bar: 5 $\mu$m.

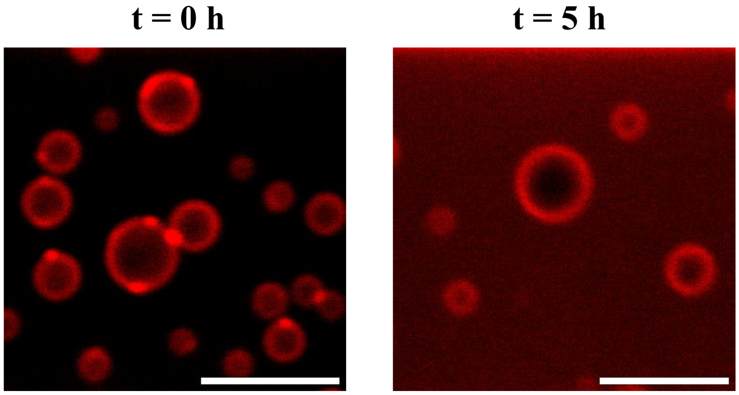


Fig. S7: Interfacial localization of designed peptide at the condensate formed by hnRNPA1-LCD (10 $\mu$M protein/peptide, 20 mM Tris (pH 7.5) and 150 mM NaCl) is consistent over 5 hours of incubation. Scale bar: 5 $\mu$m.

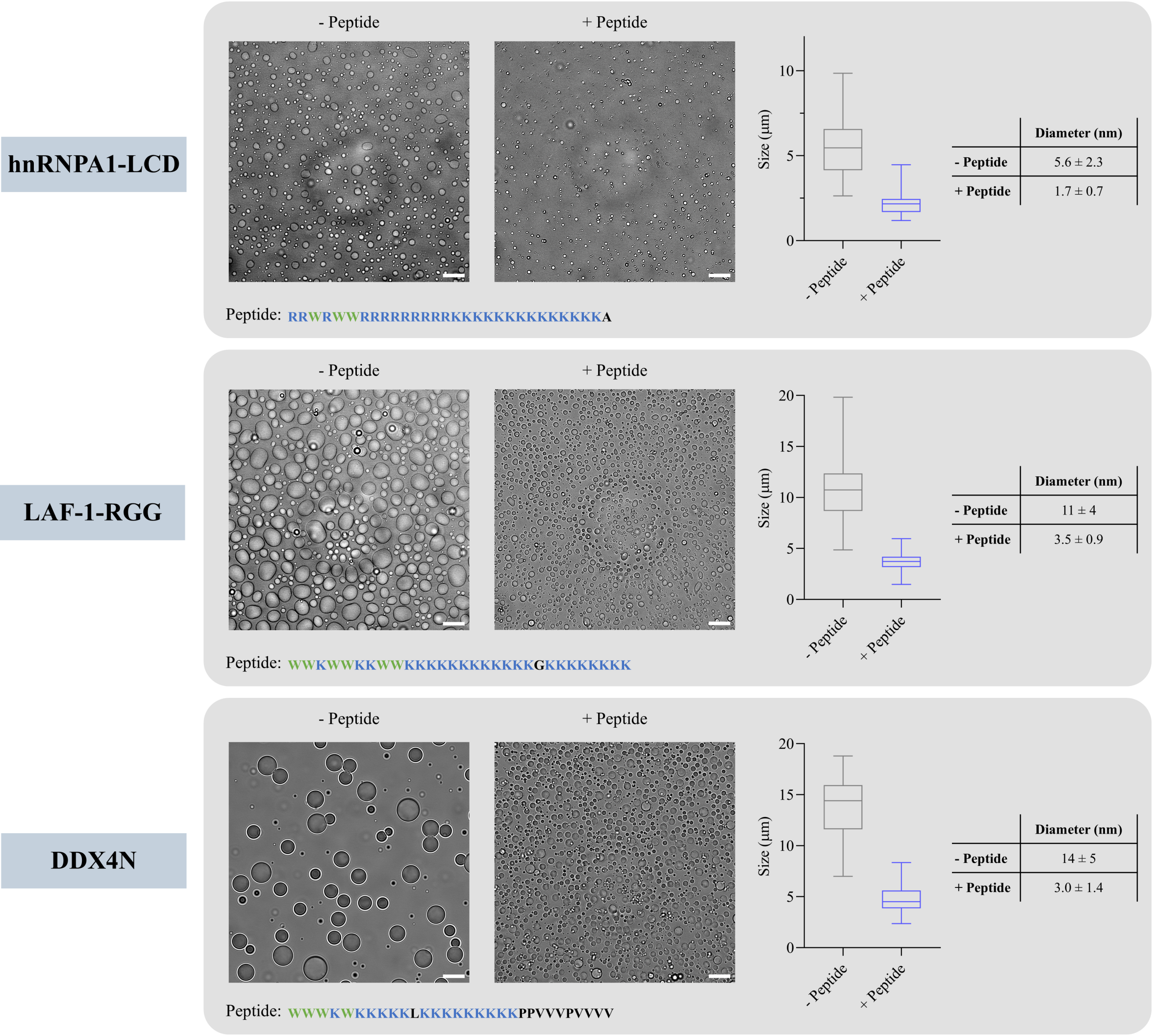


Fig. S8: Impact of interface-partitioning peptides on condensate size distribution. In all three cases, a decrease in average droplet diameter is observed. The bright-field images for hnRNPA1-LCD are identical to those in Figure 4C. The diameters reported in the table correspond to the mean ± standard deviation (n = 50). Scale bar: 20 $\mu$m.

### 6 Dynamic light scattering

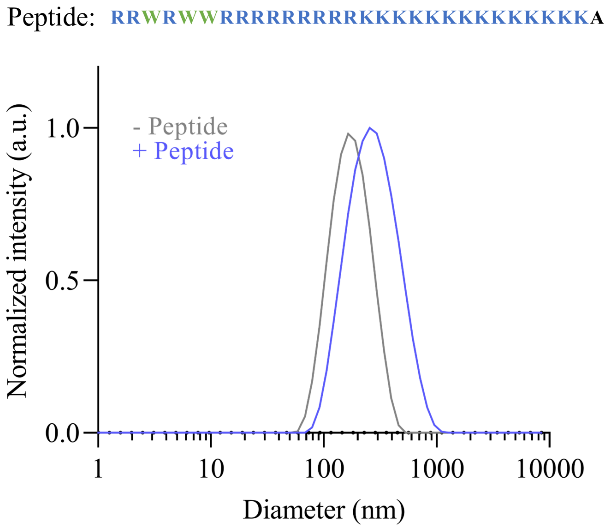


Fig. S9: Dynamic light scattering (DLS) analysis of the supernatant size distribution of hnRNPA1-LCD condensates, with and without the peptide.

### 7 Optimization results

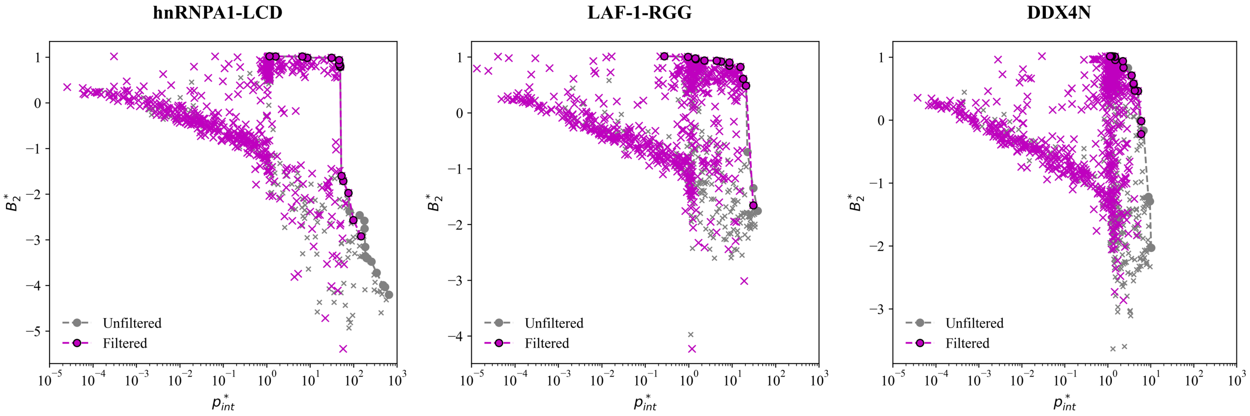


Fig. S10: Filtered optimization results for all design cases. The final filtering was performed by applying the TANGO[13] and Waltz[14] (threshold=85) aggregation predictors, as described in the main text.

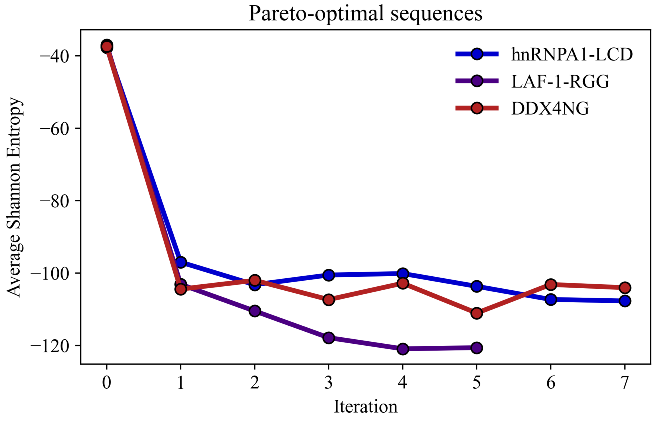


Fig. S11: Average Shannon entropy[15] of Pareto-optimal sequences across iterations for all optimization cases. The decreasing entropy indicates that the final optimal sequences were composed of only a subset of amino acids.

Tab. S1: Final Pareto front for the hnRNPA1-LCD condensate target.

| Sequence | $\Delta G_1$ [$k_\mathrm{B}T$] | $\Delta G_2$[$k_\mathrm{B}T$] | $B_2$[nm$^3$] | $p^*_{int}$ | $B^*_2$ |
|---|---|---|---|---|---|
| WWWWPWWWWPWYKKKKKKKKKKKKKKKKKK | 6.69 ± 0.15 | 12.75 ± 0.26 | -3.2E+05 ± 6.7E+04 | 6.50E+02 | -4.20 |
| WWWWPWWWYPWWKKKKKKKKKKKKKKKKKK | 6.40 ± 0.50 | 12.93 ± 0.66 | -2.2E+05 ± 3.1E+04 | 5.26E+02 | -4.03 |
| WWWPWWWWPYWWKKKKKKKKKKKKKKKKKK | 6.33 ± 0.46 | 12.63 ± 0.59 | -2.0E+05 ± 2.1E+04 | 4.74E+02 | -3.99 |
| WWWWDWEWWWWKKKKQKKKKKKKKKKKKKK | 6.51 ± 0.27 | 11.13 ± 0.17 | -1.1E+05 ± 1.8E+04 | 3.37E+02 | -3.72 |
| WWWYPYWYYPWYWYKKKKKKKKKKKKKKKK | 5.66 ± 0.46 | 12.37 ± 0.41 | -6.0E+04 ± 1.2E+04 | 2.56E+02 | -3.48 |
| WWWWPWWWWKKKWVVKKKKVKKKKKKKKKK | 6.21 ± 0.22 | 10.48 ± 0.27 | -5.1E+04 ± 1.2E+04 | 2.08E+02 | -3.41 |
| WWWWPWWWWKKKWVVKKKKKKKKKKKKKKK | 6.35 ± 0.34 | 10.31 ± 0.63 | -5.0E+04 ± 8.4E+03 | 1.97E+02 | -3.40 |
| WWWWPWWWWKKKWVVKKKKKKKKKKKKKIK | 5.99 ± 0.20 | 10.46 ± 0.54 | -4.6E+04 ± 1.0E+04 | 1.86E+02 | -3.36 |
| WWWWPWWWGWKKWKWKKKKKKKKKKKKKKK | 5.44 ± 0.32 | 11.42 ± 0.47 | -2.9E+04 ± 7.8E+03 | 1.84E+02 | -3.16 |
| WWWWPWWKWWKKWWKKKKKKKKKKKKKKKK | 5.45 ± 0.48 | 11.25 ± 0.56 | -1.1E+04 ± 1.1E+03 | 1.78E+02 | -2.75 |
| WWWWPWWKWKWKWWKKKKKKKKKKKKKKKK | 5.40 ± 0.20 | 11.32 ± 0.41 | -7.6E+03 ± 1.0E+03 | 1.75E+02 | -2.58 |
| WWWPWWWKWKKWWWKKKKKKKKKKKKKKKK | 5.55 ± 0.37 | 10.30 ± 0.51 | -5.8E+03 ± 5.0E+02 | 1.38E+02 | -2.46 |
| WWWWPWGWWKKKWWWKKKKKKKKKKKKKKK | 4.80 ± 0.26 | 10.06 ± 0.59 | -4.7E+03 ± 6.9E+02 | 7.98E+01 | -2.37 |
| RRWRWWRRRRRRRRRKKKKKKKKKKKKKKA | 4.67 ± 0.24 | 10.16 ± 0.61 | -1.9E+03 ± 5.7E+02 | 7.56E+01 | -1.97 |
| WRWWRRRRRRRRRRRTKKKKKKKKKKKKKK | 4.48 ± 0.45 | 9.70 ± 0.66 | -1.0E+03 ± 2.5E+02 | 5.72E+01 | -1.71 |
| RWRRWWRRRRRRRRRKKKKKKKKKKKKKKA | 4.03 ± 0.19 | 11.16 ± 0.45 | -7.8E+02 ± 1.2E+02 | 5.21E+01 | -1.60 |
| WWRRRRRRRRRRRRRKKKKKKKKKKKKKKA | 4.10 ± 0.63 | 10.19 ± 0.65 | 1.0E+02 ± 6.1E+00 | 4.90E+01 | 0.79 |
| RRRRWWRRRRRRRRRKKKKKKKKKKKKAKA | 4.26 ± 0.43 | 9.67 ± 0.43 | 1.1E+02 ± 8.4E+00 | 4.88E+01 | 0.81 |
| WRWRRRRRRRRRRRRRKKKKKKKKKKKKKA | 3.92 ± 0.24 | 10.95 ± 0.34 | 1.2E+02 ± 2.5E+00 | 4.62E+01 | 0.85 |
| WRRRRRWRRRRRRRRKKKKTKKKKKKKKKA | 4.41 ± 0.50 | 9.25 ± 0.74 | 1.5E+02 ± 4.0E+00 | 4.61E+01 | 0.94 |
| RRRRRRRRRRRRRRRRKKKKKKKKKKKKAA | 3.69 ± 0.79 | 9.49 ± 1.43 | 1.7E+02 ± 2.2E+00 | 3.08E+01 | 0.99 |
| RKRRKRRRRRRRRRRRRRKKKKKKKKKKAA | 2.42 ± 0.47 | 8.19 ± 0.78 | 1.7E+02 ± 2.0E+00 | 8.54E+00 | 0.99 |
| RRRRRKKRKRRRRRRRRRRKKKKKKKKKKK | 2.42 ± 0.30 | 7.35 ± 0.45 | 1.9E+02 ± 3.6E+00 | 6.52E+00 | 1.02 |
| RRRRRKKKKKKRRRRRRRRKRRRKKKKKKK | 0.67 ± 0.44 | 6.74 ± 0.62 | 1.9E+02 ± 1.7E+00 | 1.59E+00 | 1.02 |
| RRRRRRRRRRRRRRRRRRRRRRRRRRRRRR | 0.15 ± 0.16 | 35.50 ± 0.71 | 1.9E+02 ± 2.4E+00 | 1.16E+00 | 1.02 |

Tab. S2: Final Pareto front for the hnRNPA1-LCD condensate target, after applying TANGO[13] and Waltz[14] (threshold=85) aggregation filters.

| Sequence | $\Delta G_1$ [$k_\mathrm{B}T$] | $\Delta G_2$[$k_\mathrm{B}T$] | $B_2$[nm$^3$] | $p^*_{int}$ | $B^*_2$ |
|---|---|---|---|---|---|
| WWWWKKKWWWWPWWKKKKKKKKKKKKKKKK | 5.60 ± 0.24 | 10.41 ± 0.45 | -1.7E+04 ± 2.3E+03 | 1.49E+02 | -2.92 |
| WWWWKKKWWWWPWKWKKKKKKKKKKKKKKK | 5.36 ± 0.51 | 9.82 ± 0.47 | -7.4E+03 ± 1.2E+03 | 9.86E+01 | -2.57 |
| RRWRWWRRRRRRRRRKKKKKKKKKKKKKKA | 4.67 ± 0.24 | 10.16 ± 0.61 | -1.9E+03 ± 5.7E+02 | 7.56E+01 | -1.97 |
| WRWWRRRRRRRRRRRTKKKKKKKKKKKKKK | 4.48 ± 0.45 | 9.70 ± 0.66 | -1.0E+03 ± 2.5E+02 | 5.72E+01 | -1.71 |
| RWRRWWRRRRRRRRRKKKKKKKKKKKKKKA | 4.03 ± 0.19 | 11.16 ± 0.45 | -7.8E+02 ± 1.2E+02 | 5.21E+01 | -1.60 |
| WWRRRRRRRRRRRRRKKKKKKKKKKKKKKA | 4.10 ± 0.63 | 10.19 ± 0.65 | 1.0E+02 ± 6.1E+00 | 4.90E+01 | 0.79 |
| RRRRWWRRRRRRRRRKKKKKKKKKKKKAKA | 4.26 ± 0.43 | 9.67 ± 0.43 | 1.1E+02 ± 8.4E+00 | 4.88E+01 | 0.81 |
| WRWRRRRRRRRRRRRRKKKKKKKKKKKKKA | 3.92 ± 0.24 | 10.95 ± 0.34 | 1.2E+02 ± 2.5E+00 | 4.62E+01 | 0.85 |
| WRRRRRWRRRRRRRRKKKKTKKKKKKKKKA | 4.41 ± 0.50 | 9.25 ± 0.74 | 1.5E+02 ± 4.0E+00 | 4.61E+01 | 0.94 |
| RRRRRRRRRRRRRRRRKKKKKKKKKKKKAA | 3.69 ± 0.79 | 9.49 ± 1.43 | 1.7E+02 ± 2.2E+00 | 3.08E+01 | 0.99 |
| RKRRKRRRRRRRRRRRRRKKKKKKKKKKAA | 2.42 ± 0.47 | 8.19 ± 0.78 | 1.7E+02 ± 2.0E+00 | 8.54E+00 | 0.99 |
| RRRRRKKRKRRRRRRRRRRKKKKKKKKKKK | 2.42 ± 0.30 | 7.35 ± 0.45 | 1.9E+02 ± 3.6E+00 | 6.52E+00 | 1.02 |
| RRRRRKKKKKKRRRRRRRRKRRRKKKKKKK | 0.67 ± 0.44 | 6.74 ± 0.62 | 1.9E+02 ± 1.7E+00 | 1.59E+00 | 1.02 |
| RRRRRRRRRRRRRRRRRRRRRRRRRRRRRR | 0.15 ± 0.16 | 35.50 ± 0.71 | 1.9E+02 ± 2.4E+00 | 1.16E+00 | 1.02 |

Tab. S3: Final Pareto front for the LAF-1-RGG condensate target.

| Sequence | $\Delta G_1$ [$k_\mathrm{B}T$] | $\Delta G_2 [k_\mathrm{B}T]$ | $B_2[\mathrm{nm}^3]$ | $p^*_{int}$ | $B^*_2$ |
|---|---|---|---|---|---|
| WWWWPWWKKKKKKKKKKKKKKKKKKKKKKKKK | 3.70 ± 0.48 | 11.36 ± 0.70 | -1.1E+03 ± 9.9E+01 | 3.86E+01 | -1.76 |
| WWWWPWKWKKKKKKKKKKKKKKKKKKKKKKKK | 3.47 ± 0.16 | 11.02 ± 0.33 | -4.2E+02 ± 3.5E+01 | 3.06E+01 | -1.35 |
| WWWWPKWKWKKKKKKKKKKKKKKKKKKKKKGK | 3.23 ± 0.37 | 9.92 ± 0.39 | -8.0E+01 ± 1.6E+01 | 2.25E+01 | -0.70 |
| WWWWKKKWWKKVKKVKKVKKKKKKKKKKKK | 3.48 ± 0.21 | 8.68 ± 0.41 | 4.1E+01 ± 7.6E+00 | 2.09E+01 | 0.49 |
| WWWWKKKWWKKKKCKKKKKKKKKKKKKKKK | 3.07 ± 0.57 | 9.29 ± 0.65 | 6.2E+01 ± 8.0E+00 | 1.79E+01 | 0.61 |
| WWKWWKKWWKKKKKKKKKKKGKKKKKKK | 2.99 ± 0.60 | 8.87 ± 0.66 | 1.1E+02 ± 4.4E+00 | 1.55E+01 | 0.83 |
| WWKWKKWWWKKKHKKKKKKKKKKKKKKKKK | 2.21 ± 0.54 | 9.72 ± 0.81 | 1.2E+02 ± 2.7E+00 | 8.67E+00 | 0.85 |
| WKWKKKKWWWWKKKKKKKKKKKKKKKKKKKK | 2.48 ± 0.38 | 8.06 ± 0.61 | 1.4E+02 ± 3.1E+00 | 8.65E+00 | 0.91 |
| KKKWWKWWKKWWKKKKKKKKKKKKKKKKKK | 2.04 ± 0.43 | 7.65 ± 0.71 | 1.5E+02 ± 3.5E+00 | 5.61E+00 | 0.92 |
| KKKKKWWWKWKKWWKKKKKKKKKKKKKKKK | 1.73 ± 0.39 | 7.60 ± 0.27 | 1.5E+02 ± 2.9E+00 | 4.39E+00 | 0.94 |
| KKKKKKKWWKWKWKWKWKKKKKKKKKKKK | 1.35 ± 0.55 | 6.33 ± 0.46 | 1.5E+02 ± 2.3E+00 | 2.29E+00 | 0.94 |
| WWEEEEEEEEEEEEEEEDEEEEEEEEDDDDD | 0.46 ± 0.25 | 7.51 ± 0.30 | 1.6E+02 ± 1.7E+00 | 1.46E+00 | 0.96 |
| WEEEEEEEEEEEEEEEEEEEEEEEWEEEEE | 0.54 ± 0.30 | 6.71 ± 0.35 | 1.7E+02 ± 1.9E+00 | 1.42E+00 | 0.98 |
| RRRRRRRRRRRRRRKKKKKKRKRRRKKKYK | 0.13 ± 0.15 | 6.41 ± 0.58 | 1.8E+02 ± 1.4E+00 | 9.61E-01 | 1.00 |
| RRRRRRRRKKKKKKKKKKRRRRRRRRRKKK | 0.21 ± 0.18 | 3.55 ± 0.55 | 1.9E+02 ± 3.1E+00 | 2.72E-01 | 1.02 |

Tab. S4: Final Pareto front for the LAF-1-RGG condensate target, after applying TANGO[13] and Waltz[14] (threshold=85) aggregation filters.

| Sequence | $\Delta G_1$ [$k_\mathrm{B}T$] | $\Delta G_2 [k_\mathrm{B}T]$ | $B_2[\mathrm{nm}^3]$ | $p^*_{int}$ | $B^*_2$ |
|---|---|---|---|---|---|
| WWWPWWWKKKKKKKKKKKKKKKKSKKKGKKK | 3.46 ± 0.48 | 11.28 ± 0.43 | -8.9E+02 ± 1.3E+02 | 3.05E+01 | -1.66 |
| WWWWKKKWWKKVKKVKKVKKKKKKKKKKKK | 3.48 ± 0.21 | 8.68 ± 0.41 | 4.1E+01 ± 7.6E+00 | 2.09E+01 | 0.49 |
| WWWWKKKWWKKKKCKKKKKKKKKKKKKKKK | 3.07 ± 0.57 | 9.29 ± 0.65 | 6.2E+01 ± 8.0E+00 | 1.79E+01 | 0.61 |
| WWKWWKKWWKKKKKKKKKKKGKKKKKKK | 2.99 ± 0.60 | 8.87 ± 0.66 | 1.1E+02 ± 4.4E+00 | 1.55E+01 | 0.83 |
| WWKWKKWWWKKKHKKKKKKKKKKKKKKKKK | 2.21 ± 0.54 | 9.72 ± 0.81 | 1.2E+02 ± 2.7E+00 | 8.67E+00 | 0.85 |
| WKWKKKKWWWWKKKKKKKKKKKKKKKKKKKK | 2.48 ± 0.38 | 8.06 ± 0.61 | 1.4E+02 ± 3.1E+00 | 8.65E+00 | 0.91 |
| KKKWWKWWKKWWKKKKKKKKKKKKKKKKKK | 2.04 ± 0.43 | 7.65 ± 0.71 | 1.5E+02 ± 3.5E+00 | 5.61E+00 | 0.92 |
| KKKKKWWWKWKKWWKKKKKKKKKKKKKKKK | 1.73 ± 0.39 | 7.60 ± 0.27 | 1.5E+02 ± 2.9E+00 | 4.39E+00 | 0.94 |
| KKKKKKKWWKWKWKWKWKKKKKKKKKKKK | 1.35 ± 0.55 | 6.33 ± 0.46 | 1.5E+02 ± 2.3E+00 | 2.29E+00 | 0.94 |
| WWEEEEEEEEEEEEEEEDEEEEEEEEDDDDD | 0.46 ± 0.25 | 7.51 ± 0.30 | 1.6E+02 ± 1.7E+00 | 1.46E+00 | 0.96 |
| WEEEEEEEEEEEEEEEEEEEEEEEWEEEEE | 0.54 ± 0.30 | 6.71 ± 0.35 | 1.7E+02 ± 1.9E+00 | 1.42E+00 | 0.98 |
| RRRRRRRRRRRRRRKKKKKKRKRRRKKKYK | 0.13 ± 0.15 | 6.41 ± 0.58 | 1.8E+02 ± 1.4E+00 | 9.61E-01 | 1.00 |
| RRRRRRRRKKKKKKKKKKRRRRRRRRRKKK | 0.21 ± 0.18 | 3.55 ± 0.55 | 1.9E+02 ± 3.1E+00 | 2.72E-01 | 1.02 |

Tab. S5: Final Pareto front for the DDX4N condensate target.

| Sequence | $\Delta G_1$ [$k_\mathrm{B}T$] | $\Delta G_2[k_\mathrm{B}T]$ | $B_2[\mathrm{nm}^3]$ | $p^*_{int}$ | $B^*_2$ |
|---|---|---|---|---|---|
| WWWWPWFVKKKKVVVPVVVVPVVVVPVVVV | 2.48 ± 0.39 | 8.89 ± 0.43 | -2.1E+03 ± 2.8E+02 | 1.02E+01 | -2.03 |
| WWWWPWKKKKKKKKKPVVVVPVVVVPVVVV | 2.36 ± 0.37 | 9.24 ± 0.29 | -3.7E+02 ± 4.5E+01 | 9.56E+00 | -1.29 |
| WWWWPWKKKKKKVKKVKKVVPVVVVPVVVV | 2.26 ± 0.44 | 9.28 ± 0.53 | -3.1E+02 ± 3.6E+01 | 8.81E+00 | -1.22 |
| WWWWKKKWKKKKKVKKKKVVPVVVVPVVVV | 2.05 ± 0.41 | 8.63 ± 0.64 | -9.2E+00 ± 5.8E+00 | 6.82E+00 | -0.17 |
| WWWWKKKFKKKKKKKKKVKVPVVVVPVVVV | 1.96 ± 0.25 | 8.27 ± 0.50 | -6.4E-01 ± 5.1E+00 | 5.99E+00 | -0.01 |
| WWWWKKKKWKKKKKKKKKVVPVVVVPVVVS | 1.77 ± 0.23 | 9.18 ± 0.33 | -1.4E-01 ± 7.0E+00 | 5.56E+00 | 0.00 |
| WWWWKKKKKKKKKKKKKKKKPPVVPVVVV | 1.68 ± 0.27 | 8.96 ± 0.44 | 3.8E+01 ± 4.3E+00 | 5.03E+00 | 0.46 |
| WWWWKKKKKKKKKKKKKKKKKVVPVVVSSS | 1.50 ± 0.39 | 9.16 ± 0.57 | 3.9E+01 ± 3.8E+00 | 4.26E+00 | 0.47 |
| WWWKWKKKKKLKKKKKKKKPPVVVPVVVV | 1.80 ± 0.42 | 7.07 ± 0.32 | 5.4E+01 ± 2.9E+00 | 3.99E+00 | 0.57 |
| WWWWKKKKKKKKKKKKKKPVVVVPVVKVKK | 1.43 ± 0.36 | 9.01 ± 0.58 | 5.6E+01 ± 4.3E+00 | 3.97E+00 | 0.58 |
| WWYWKKKKKKKKKKKKKKPPVVPVVVKKKK | 1.47 ± 0.27 | 7.58 ± 0.43 | 8.2E+01 ± 3.2E+00 | 3.56E+00 | 0.71 |
| DDDDDDDDDWDWWWDFFDDDDDDDDDDDGG | 1.09 ± 0.23 | 11.77 ± 0.63 | 1.1E+02 ± 1.8E+00 | 2.96E+00 | 0.83 |
| DDDDDDDDDDDDWWWWDDDDDDDDDGGGGG | 0.87 ± 0.33 | 9.14 ± 0.50 | 1.2E+02 ± 2.6E+00 | 2.33E+00 | 0.83 |
| DDDDDDDDDWWDWWDDDDDDDDDDDDDDDG | 0.83 ± 0.42 | 9.25 ± 0.55 | 1.5E+02 ± 2.1E+00 | 2.24E+00 | 0.94 |
| EDDDDEDDDDDDDDDDDDYWDYWDDDDDDD | 0.46 ± 0.42 | 7.79 ± 0.94 | 1.6E+02 ± 3.4E+00 | 1.48E+00 | 0.96 |
| KKKKKKKKHKRKKKRRRRRRKKKKKKKKKK | 0.48 ± 0.32 | 7.42 ± 0.44 | 1.8E+02 ± 2.2E+00 | 1.47E+00 | 1.01 |
| RRRRRKKKKKKKKKKKKRRRRRRRRKKKKK | 0.23 ± 0.29 | 13.34 ± 0.54 | 1.9E+02 ± 4.0E+00 | 1.26E+00 | 1.02 |
| RRRRKKKKKKKKKKKRRRRRRRRRRRRKRK | 0.20 ± 0.25 | 17.50 ± 0.77 | 1.9E+02 ± 2.2E+00 | 1.22E+00 | 1.02 |
| RRRRRRRKKKKKKKKKKKKKRKRKRRRRRR | 0.18 ± 0.13 | 15.48 ± 0.32 | 1.9E+02 ± 2.3E+00 | 1.19E+00 | 1.02 |
| RRRRRRRKKKKKKKKKKKKKKKRKRRKRKK | 0.13 ± 0.08 | 10.44 ± 0.42 | 1.9E+02 ± 2.0E+00 | 1.13E+00 | 1.02 |

Tab. S6: Final Pareto front for the DDX4N condensate target, after applying TANGO[13] and Waltz[14] (threshold=85) aggregation filters.

| Sequence | $\Delta G_1$ [$k_\mathrm{B}T$] | $\Delta G_2[k_\mathrm{B}T]$ | $B_2[\mathrm{nm}^3]$ | $p^*_{int}$ | $B^*_2$ |
|---|---|---|---|---|---|
| WWWWKKKWKKVKVKKKVKVVPKVVVPVVVV | 2.01 ± 0.42 | 8.04 ± 0.51 | -1.3E+01 ± 6.3E+00 | 6.01E+00 | -0.22 |
| WWWWKKKFKKKKKKKKKVKVPVVVVPVVVV | 1.96 ± 0.25 | 8.27 ± 0.50 | -6.4E-01 ± 5.1E+00 | 5.99E+00 | -0.01 |
| WWWWKKKKKKKKKKKKKKKKPPVVPVVVV | 1.68 ± 0.27 | 8.96 ± 0.44 | 3.8E+01 ± 4.3E+00 | 5.03E+00 | 0.46 |
| WWWWKKKKKKKKKKKKKKKKKVVPVVVSSS | 1.50 ± 0.39 | 9.16 ± 0.57 | 3.9E+01 ± 3.8E+00 | 4.26E+00 | 0.47 |
| WWWKWKKKKKLKKKKKKKKPPVVVPVVVV | 1.80 ± 0.42 | 7.07 ± 0.32 | 5.4E+01 ± 2.9E+00 | 3.99E+00 | 0.57 |
| WWWWKKKKKKKKKKKKKKPVVVVPVVKVKK | 1.43 ± 0.36 | 9.01 ± 0.58 | 5.6E+01 ± 4.3E+00 | 3.97E+00 | 0.58 |
| WWYWKKKKKKKKKKKKKKPPVVPVVVKKKK | 1.47 ± 0.27 | 7.58 ± 0.43 | 8.2E+01 ± 3.2E+00 | 3.56E+00 | 0.71 |
| DDDDDDDDDDDDWWWWDDDDDDDDDGGGGG | 0.87 ± 0.33 | 9.14 ± 0.50 | 1.2E+02 ± 2.6E+00 | 2.33E+00 | 0.83 |
| DDDDDDDDDWWDWWDDDDDDDDDDDDDDDG | 0.83 ± 0.42 | 9.25 ± 0.55 | 1.5E+02 ± 2.1E+00 | 2.24E+00 | 0.94 |
| EDDDDEDDDDDDDDDDDDYWDYWDDDDDDD | 0.46 ± 0.42 | 7.79 ± 0.94 | 1.6E+02 ± 3.4E+00 | 1.48E+00 | 0.96 |
| KKKKKKKKHKRKKKRRRRRRKKKKKKKKKK | 0.48 ± 0.32 | 7.42 ± 0.44 | 1.8E+02 ± 2.2E+00 | 1.47E+00 | 1.01 |
| RRRRRKKKKKKKKKKKKRRRRRRRRKKKKK | 0.23 ± 0.29 | 13.34 ± 0.54 | 1.9E+02 ± 4.0E+00 | 1.26E+00 | 1.02 |
| RRRRKKKKKKKKKKKRRRRRRRRRRRRKRK | 0.20 ± 0.25 | 17.50 ± 0.77 | 1.9E+02 ± 2.2E+00 | 1.22E+00 | 1.02 |
| RRRRRRRKKKKKKKKKKKKKRKRKRRRRRR | 0.18 ± 0.13 | 15.48 ± 0.32 | 1.9E+02 ± 2.3E+00 | 1.19E+00 | 1.02 |
| RRRRRRRKKKKKKKKKKKKKKKRKRRKRKK | 0.13 ± 0.08 | 10.44 ± 0.42 | 1.9E+02 ± 2.0E+00 | 1.13E+00 | 1.02 |
